# Inferring high-dimensional pathways of trait acquisition in evolution and disease

**DOI:** 10.1101/409656

**Authors:** Sam F. Greenbury, Mauricio Barahona, Iain G. Johnston

## Abstract

The explosion of data throughout the sciences provides unprecedented opportunities to learn about the dynamics of evolution and disease progression. Here, we describe a highly generalisable statistical platform to infer the dynamic pathways by which many, potentially interacting, discrete traits are acquired or lost over time in biological processes. The platform uses HyperTraPS (hypercubic transition path sampling) to learn progression pathways from cross-sectional, longitudinal, or phylogenetically-linked data with unprecedented efficiency, readily distinguishing multiple competing pathways, and identifying the most parsimonious mechanisms underlying given observations. Its Bayesian structure quantifies uncertainty in pathway structure and allows interpretable predictions of behaviours, such as which symptom a patient will acquire next. We exploit the model’s topology to provide visualisation tools for intuitive assessment of multiple, variable pathways. We apply the method to ovarian cancer progression and the evolution of multidrug resistance in tuberculosis, demonstrating its power to reveal previously undetected dynamic pathways.

## 1. Introduction

Many problems in biomedicine and throughout the sciences can be thought of as the serial stochastic acquisition of discrete features or traits. These traits may be, for example, the symptoms experienced by a patient during progressive diseases, the genetic and physiological features underlying cancer progression, or the acquisition of drug-resistance traits in pathogens. Understanding the dynamics of these processes has the potential to inform targetted therapies, reveal biological mechanisms, and predict future behaviours, and has been an open challenge throughout the data explosion in biomedical sciences Colijn et al. (2017).

Existing methods to reconstruct the past, and predict the future, of processes involving discrete trait acquisition have emerged from both the cancer science and evolutionary literature. Authors attempt to classify cancer-related alterations into progressive ‘hallmarks’ (Hanahan and Weinberg, 2000, 2011). With the advent of high-throughput sequencing, a large amount of effort has been placed in utilising phylogenetic methods for understanding the way in which cancer progresses at the genetic level (Schwartz and Schäffer, 2017). Beerenwinkel et al. (2015) provide an excellent review of methods designed for these systems: a large array of methods have been developed for deriving progression models to devise better treatments and interventions through understanding the way and order in which alterations are acquired (Beerenwinkel et al., 2015; Schwartz and Schäffer, 2017). These range from stochastic models employing Markov chains for acquisition on graphs such as in Hjelm et al. (2006), to Bayesian network approaches where trees, forests or a directed acyclic graph (DAG) are to be inferred from the data (Szabo and Boucher, 2002; Beerenwinkel et al., 2007; Gerstung et al., 2009; Loohuis et al., 2014; Ramazzotti et al., 2015). A major branch of this field has focussed on independent samples exhibiting differing presence of alterations (*cross-sectional data*) for reconstructing *oncogenetic models* (Beerenwinkel et al., 2015) to discover cancer progression or potentially causal relationship of markers in patients.

Evolutionary and phylogenetic approaches for inferring trait dynamics, by contrast, must account for the relatedness of individuals and the possibility that a given state in a progressive system is inherited from an ancestor. Notable models that have attempted to solve this problem have included Bollback (2006) with Markov Chain Monte Carlo (MCMC) approaches to sampling from a master equation model for character mapping with Simmap. Other similar methods include Ordermutation (Youn and Simon, 2012) and the Reversible Jump MCMC (RJ-MCMC) methodology also applied to a master equation formulation (Pagel and Meade, 2006). Such approaches have been utilised for understanding the evolution of phenotypic traits in populations (Mahler et al., 2010; Watts et al., 2015). In connection with cancer progression, more recently modelling approaches have been developed with the aim of reconstructing phylogenetic models from sources such as single-cell sequencing data (Beerenwinkel et al., 2015; Ross and Markowetz, 2016; Zafar et al., 2017; Ramazzotti et al., 2017).

Challenges remain in applying these algorithms to dissect the dynamics of systems involving many, potentially coupled, traits. Existing methods may assume an absence of coupling or interactions between traits or be restricted to a limited number. Computational runtime often scales exponentially with the underlying number of features, and frequently exhibits challenging scaling with the number of observations and traits, limiting the applicability of the approaches to many forms of biomedical data, particular given modern trends of increasing data volumes and heterogeneity. A recent approach, HyperTraPS (hypercubic transition path sampling) (Johnston and Williams, 2016), aimed to address these issues, allowing the inference of the dynamics of many coupled traits from data following arbitrary (but known) phylogenetic relationships. HyperTraPS represents progressive dynamics as paths on a hypercubic space connecting all possible patterns of trait presence and absence, and uses observations of intermediate states to learn the most likely pathways of progress through this space. In this way, snapshot data can be used to learn the probabilistic structure of dynamic pathways, which have in turn been used to identify the mechanisms underlying the evolutionary dynamics of *L* = 65 mtDNA genes (Johnston and Williams, 2016) and *C*_3_ to *C*_4_ photosynthesis (Williams et al., 2013).

To date, HyperTraPS has only been used to address these specific evolutionary questions. However, in the current era of large-scale scientific and biomedical data, questions about the structure of dynamic pathways are expanding and becoming more pertinent to endeavours from evolutionary biology to precision medicine. Hypercubic inference represents a powerful new way of addressing these questions, but a general platform for its application, interpretation, and visualisation remains absent. Such a platform would provide many advantages over the current state of the art: large-scale datasets can be readily analysed, different types of observational data can be used (cross-sectional, longitudinal, and/or phylogenetically coupled observations); Bayesian quantification of uncertainty and a completely unrestricted set of states and transitions can be applied, and competing pathways and their detailed structure can be resolved and characterised, facilitating the identification of progression mechanisms. In principle, any dataset where the relationship of the samples is known or can be inferred is amenable to this detailed analytic approach.

Here, we address this target, presenting a novel and expansive set of developments to allow the inference of dynamic pathways from highly general datasets. We embed HyperTraPS in a new and efficient platform for parametric inference and model selection, simultaneously allowing Bayesian inference of dynamic pathways and the identification of model structures that best describe the dynamics and interactions contained within a given set of observations. This model selection guards against overfitting and reveals the extent to which interactions between features dictate the dynamics of the observed system. Models identified in this way have the strongest power to predict out-of-sample observations, which we demonstrate with synthetic and real-world examples, illustrating the predictive power of the approach. To further facilitate interpretation of the inference outcomes, we introduce approaches for intuitively visualising and comparing the high-dimensional pathways inferred from complex datasets, which may include multiple distinct orderings for the acquired traits.

We illustrate the performance of these methods in three different scenarios: with illustrative synthetic datasets; with a well-studied dataset on the progressive acquisition of genetic alterations in ovarian cancer; and with a recent large-scale dataset on drug-resistant tuberculosis. We compare this novel HyperTraPS platform to other approaches from the disease progression and evolutionary literatures for trait inference, highlighting its intersection between these fields and consequent general power and applicability. We conclude by discussing the breadth of applications in the expanding fields of precision medicine, data science, and evolutionary inference, and provide an open source package for the code.

## 2. Results

### 2.1. Inferring dynamic pathways involving coupled traits on general state spaces

HyperTraPS represents every possible state of a system with *L* features or traits (we use these terms synonymously here) as a binary string of length *L*, where 0 and 1 at the *i*^th^ position correspond respectively to absence or presence of the *i*th trait. HyperTraPS assumes that systems start with no traits present (*O*, the string of all 0s). Traits are then acquired stochastically and irreversibly, according to transition probabilities linking states. We assume that traits are acquired one-at-a-time, so that the states and transitions are embedded on a hypercubic transition graph (Fig. 1). We then consider instances of an evolving or progressing system as an ensemble of random walkers on this graph. As in a hidden Markov model Murphy (2012), observations are assumed to arise through randomly emitted signals by these walkers; a signal corresponds to the current set of acquired traits of the random walker. The task at the core of HyperTraPS is to compute the likelihood of observing a set of emissions that match the states in a dataset, given a parameterisation *W* describing the transition probabilities on the edges of the hypercube.

**Fig. 1:**
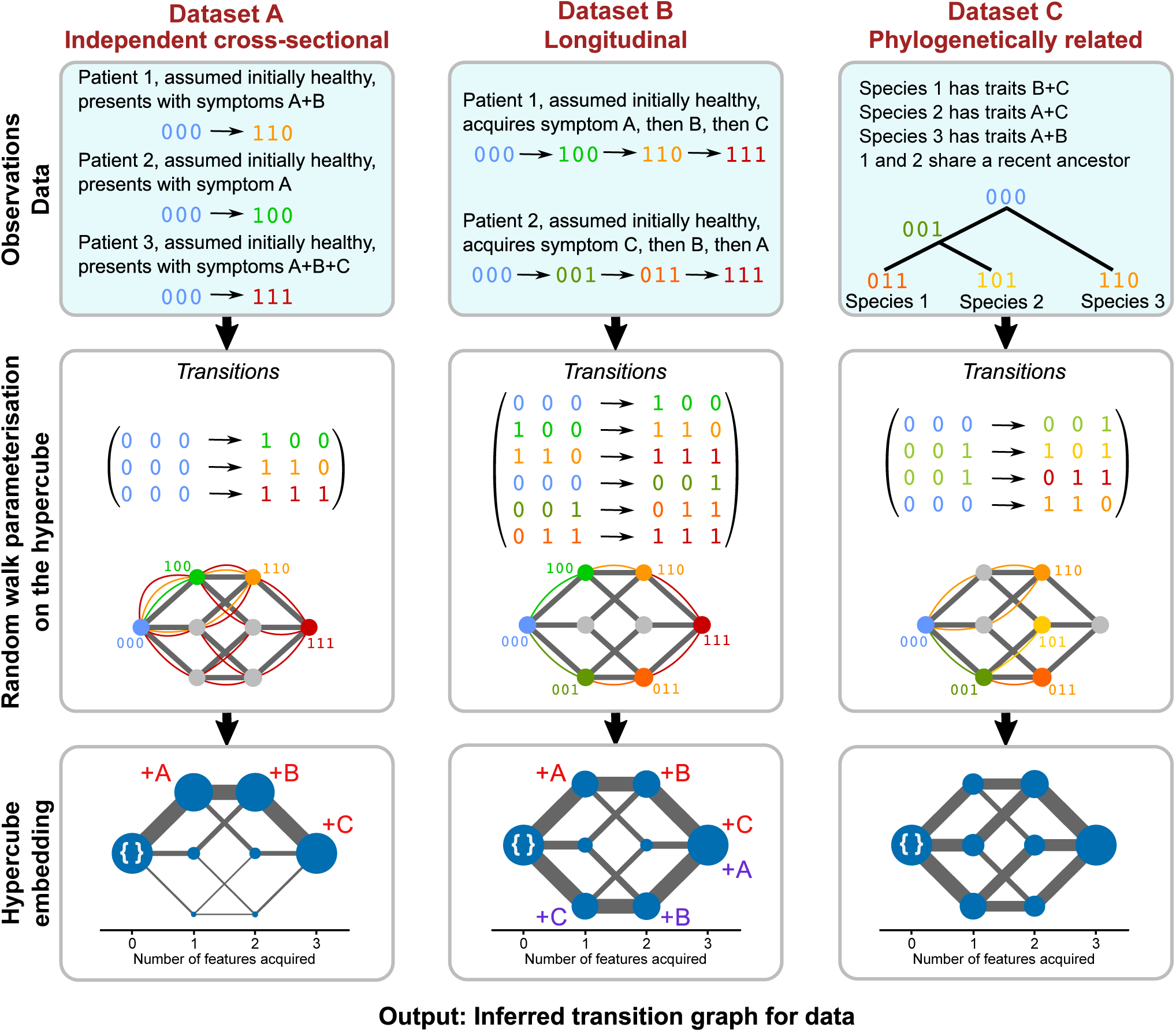
HyperTraPS allows dynamic inference with three classes of input data A, B and C. In each case, presence/absence of traits are labelled with a binary marker, and temporal relationships between observations (if present) are invoked to represent *observed samples* as *observed transitions*, yielding *transition sets*. With a dataset of transitions, the likelihood that a given set of edge weights on the underlying hypercubic transition network will give rise to the observed transitions can be calculated efficiently using a path sampling approach where *random walkers sample parameterisations*. Each (hyper)cube corresponds to the datasets (A, B and C), with colour coded curved edges and states showing the possible paths that can be taken to reach observed samples from the source state. Edge weights that reproduce the observed transitions are more likely to represent the underlying transition process. Embedding this likelihood calculation in a Bayesian inference scheme allows posterior weights on *inferred transition graphs* to be computed, constituting a complete characterisation of the dynamic systems. Edge widths and vertex areas are proportional to the posterior weighting.

In STAR Methods, we outline the HyperTraPS algorithm to estimate this likelihood given a set of observations. In contrast to previous approaches (Johnston and Williams, 2016; Williams et al., 2013), we embed the core likelihood calculation in an auxilary pseudo-marginal MCMC (APM MCMC) framework (Murray and Graham, 2015) to allow more efficient Bayesian inference of the hypercubic transition network supporting the observed dynamics. The APM MCMC embedding overcomes potential issues arising from uncertainty in the likelihood estimates for long pathway calculations (STAR Methods), better guaranteeing that the MCMC process will mix well and converge to a consistent posterior in the case of large, sparse inference challenges. APM MCMC makes it possible to address systems involving dozens of sparsely sampled traits, as we demonstrate below.

The next important consideration in this inference process is how this transition network is parameterised. Individually parameterising each of *L*2^*L-*1^ hypercubic edges represents a huge inference challenge for (likely) very little model fit reward. Instead, we propose a hierarchy of parameter representations (Fig. 6). For the *zero order model every feature has equal probability of acquisition*. *All edges on the transition network thus have the same weight, requiring no parameters*. *In the* first order model, every feature has an independent acquisition probability regardless of current state. Transition edge weights between two states are thus exclusively determined by the trait that distinguished the two states (requiring *k* = *L* parameters). In the *second order* model, every feature’s acquisition probability depends independently on the presence of each other feature. Transition edge weights between two states thus depend on the distinguishing trait and the presence/absence of each other trait (requiring *L*^2^ parameters; as in (Johnston and Williams, 2016)). Higher order models, including the full *L*2^*L-*1^ set naturally follow, introducing more complex interactions between the co-occurrence of features (as in (Williams et al., 2013)). The appropriate choice of parameterisation is dictated by the generative processes underlying the observed data; if trait acquisitions are independent, the parsimonious first-order model is more appropriate; if traits interact pairwise, the second-order model will be required to capture the dynamics. A given dataset may be best described by an intermediate representation between two of these cases.

To identify the optimal parameter representation for a given dataset, we introduce methods for regularising the inferred model parameterisations (see STAR Methods), allowing the appropriate choice of model structure to describe the observed data and a means of generating maximum likelihood parameterisations without overfitting. The regularisation allows us to distinguish simple cases, where all dynamics can be described by traits behaving independently, from more complex cases where the acquisition of one or more traits influences the probability of acquisition of other traits. This combination of an efficient and general inference platform, a process for model selection, and a new toolbox for visualising and interpreting inferred posteriors, allows us for the first time to apply HyperTraPS to a dramatically expanded range of biomedical questions.

In order to illustrate the ability of HyperTraPS to characterise dynamics from independent cross-sectional samples, we constructed three cross-sectional datasets with different underlying progressions. First, 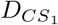 involves samples taken uniformly from each state along a single trajectory, where features are accumulated from *left to right*. For example, for *L* = 3, the sequence of acquisition is 000*→*100*→*110*→*111. Second, 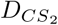 involves samples taken uniformly from states along two distinct progression pathways with exactly opposing temporal ordering of acquisition: one where features are acquired from *left to right* and the other where features are acquired from *right to left*. For example, for *L* = 3, this corresponds to the two trajectories 000 *→* 100 *→* 110 *→* 111 and 000 *→* 001 *→* 011 *→* 111. Lastly, a composite of 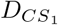 and 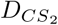 such that 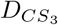 is the linear combination: 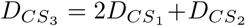. In this case, we have a dominant progression underlying the dataset but with a significant alternative progression also being present.

We chose these structures to illustrate HyperTraPS’s ability to infer both *single* 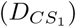, *multiple competing* 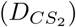 pathways and *multiple differentially weighted competing* 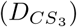 pathways. For the single pathway, traits can be independent – a suitable ordering of the ‘basal rates’ is sufficient to generate 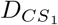. By contrast, competing pathways require traits to interact – acquisition of traits on one pathway in 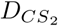 and 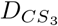 must repress acquisition of traits on the other pathway. Importantly, the samples used in the dataset are cross-sectional and therefore the underlying progressions that generated the samples is to be inferred; the data is input simply as the set of transitions from the state with no features present to the observed signals.

In Fig. 2, we demonstrate structure of the data, pathway inference, model regularisation, and validation for each of the 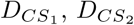 and 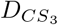. Fig. 2A(i)-(iii) shows the visual structure of the data with samples plotted against features with black rectangles indicating the presence of a feature in the sample. To visualise the inferred dynamic behaviour, we use a custom algorithm (described in further detail in STAR methods) to project the inferred hypercubic transition network into two dimensions, arranging states with increasing numbers of features from left to right (Fig. 2B). A single dominant progression is clear for Fig. 2B(i), while the two progressions are clearly shown in Fig. 2B(ii) and 2B(iii). Fig. 2C(i)-(iii) show an alternative representation: the posterior probabilities with which each trait is acquired in each possible ordering. Again, the dynamics corresponding to the simple single pathway and the more complex competing two-pathway model are clearly visible.

**Fig. 2:**
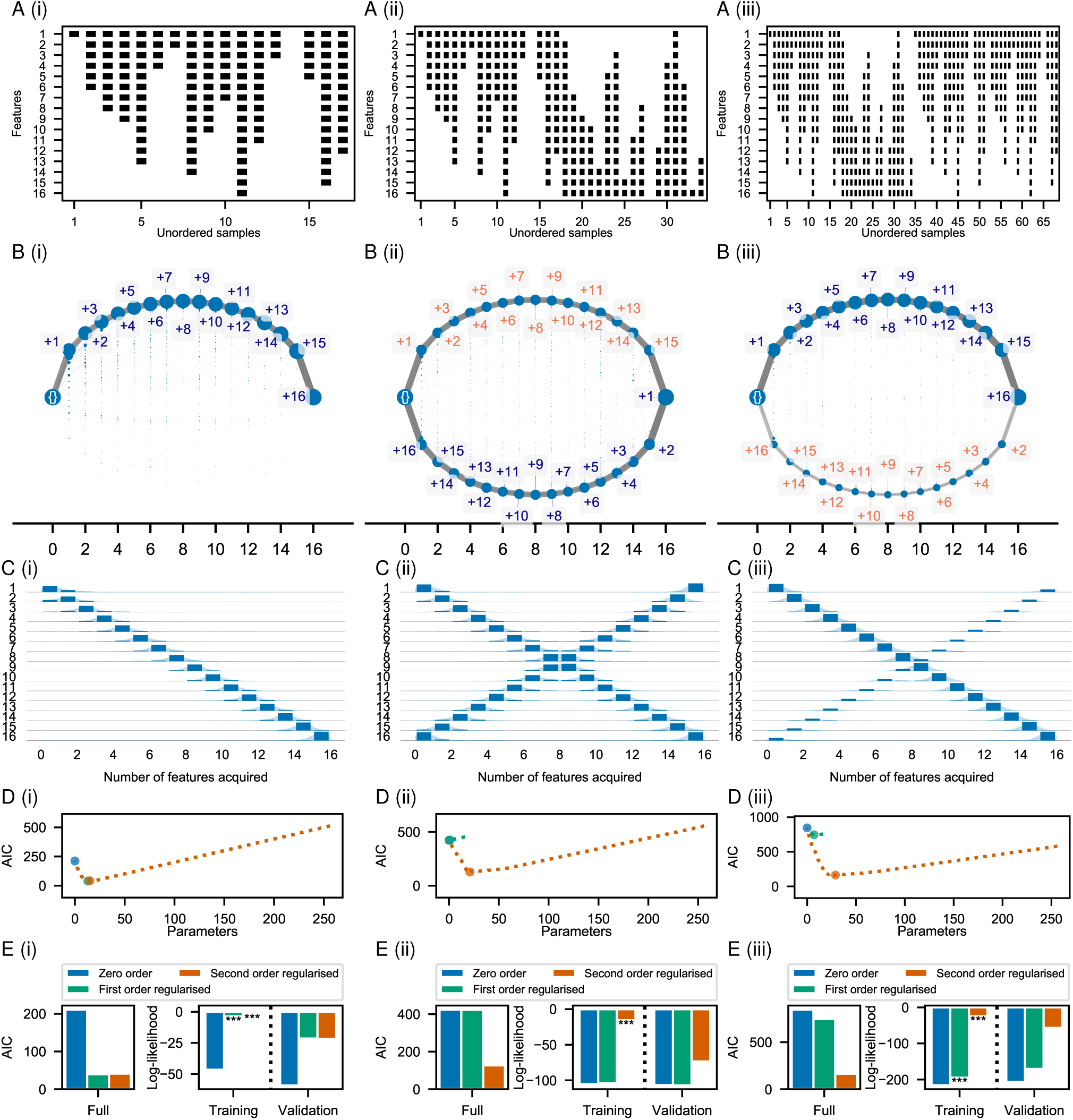
Inference, regularisation, and validation of HyperTraPS platform for two synthetic datasets. Synthetic dataset 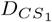 (i) supports only a single pathway; 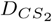 (ii) supports two competing pathways; and 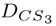 (iii) supports competing pathways but with a single dominant pathway. A. The cross-sectional datasets are represented with a trait being present in a sample where a black box is present in the corresponding feature row. The samples are unordered to highlight the platform’s ability to infer progressions from cross-sectional data. B. Inferred paths across hypercube. Random walks from the empty state “{}” state to the “all features state” are performed across parameterisations from the posterior for each dataset. Edge widths and node areas are proportional to the number of times edges/nodes are encountered. States are plotted from left to right in order of the number of features acquired in the state using the embedding and labelling procedure described in STAR methods. The single pathway clearly dominates in (i), while the two pathways present in 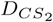 and 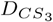 are clearly observable in (ii) and (iii). C. Inferred ordering of acquisitions. Blue bars give the probability that a feature (horizontal axis) is acquired at a given step (vertical axis). Bimodality in ordering posteriors ((ii) and (iii)) and reflect the presence distinct progressions that exist in the underlying dynamics. D. Regularisation of the inferred models. The dashed line shows the AIC score as parameters are greedily pruned from approximate first- and second-order maximum likelihood parameterisations. Each model and its corresponding AIC score is shown in the plot. The turning point illustrating an optimal sparser parameterisation denoted as the first- and second-order regularised models respectively. *Simpler dataset (i) requires fewer parameters than the more complex datasets (ii) and (iii)*. E. For each regularised model type, the AIC of the full dataset, and log-likelihoods of the training dataset and test dataset (as described in the text). For the full and training dataset, stars give the *p*-value from a likelihood ratio in comparison to the zero order model (the null model) with significance levels of *\*\*\*<* 0.001, *\*\*<* 0.01 and *\*<* 0.05. The first order regularised model is sufficient for the single pathway, while the second order regularised model is necessary for (ii) and (iii).

In this extreme example, the inferred ordering distributions for all but the central traits in 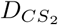 and 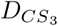 exhibit *bimodality*. Generally in such histograms from HyperTraPS posteriors, bimodality (and multimodality more generally) reflects structurally distinct progression pathways (for example, where a feature can be acquired early or late, but not at intermediate stages), while unimodal distributions reflect sets of pathways with a consistent structural trend. The width of such modes reflects the amount of noise and variation in the order for which a feature is acquired in the progression associated with the mode. Multimodal distributions in these plots provide a suggestive signature of distinct dynamic pathways of the system. In Appendix D.1, we compare this inference of competing pathways to existing alternative approaches and show that HyperTraPS has a unique ability to resolve and characterise multiple progressive pathways.

To highlight the potential for HyperTraPS to identify, disambiguate and characterise competing pathways, we applied an alternative method, Capri (Ramazzotti et al., 2015), to these synthetic datasets. Capri attempts to produce a maximum-likelihood directed acyclic graph (DAG) reflecting causal relationships between traits, given inference of causal relationships from data. For the single-pathway case 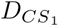, Capri and HyperTraPS yield tightly comparable results (Fig. D.12B): in both cases, the temporal relationship (first to last trait acquisitions) is captured. However, for the two-pathway case 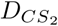 and 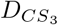, the DAG inferred by Capri does not directly reflect the dynamics of the system, which are captured by HyperTraPS (Fig. D.12A(ii)-(iii) and Fig. D.12B(ii)-(iii)). Instead, the DAG inference is rather confounded by the competing pathways and reports an indirect mix of causalities rather than the clearly separated pathways inferred by HyperTraPS.

FIG. 2D demonstrates that the model regularisation process (described in STAR Methods) dramatically reduces extraneous parameters at negligible cost to model fit. The optimal parameterisation for 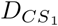 contains around *L* terms, as illustrated by the overlapping points for the first order and second order regularised models in Fig. 2D(i). The interaction terms included in the second order regularised model provide additional explanatory power for 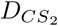 and 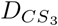, with the regularised second order model having a considerably better AIC score than the first order regularised model. This improvement is a consequence of the trait-interaction terms in the second-order model allowing the required cross-repression of pathways, making it a better explanatory model in this case.

To validate these findings and explore the predictive power of this approach, we split the data into two halves to form a training and test dataset. We obtained posteriors from the training set for each model, and computed the likelihood associated with the test set for these inferred posteriors. Fig. 2E(i)-(iii) shows the AIC scores for the full model, and the log-likelihoods for training and validation datasets. For the single-path dataset, the first order model and second order model provide similar explanatory power in the full model, and predictive power in the validation experiment, both improved over the zero order model (null model). For the two-path dataset 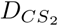, the second order model enhances predictive power compared to both the first order and null models, and regularisation improves the parsimony of this model with no cost to model fit (*p <* 0.001 for the a likelihood ratio test against the null model). For 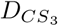, the first order regularised model has a lower AIC than the zero order model as it captures the dominant progression. However, the second order regularised model performs much better still through the ability to capture both the dominant and secondary progression present in the dataset, illustrated by lower AIC and larger log-likelihoods in the training and validation datasets.

### 2.2 Application to cross-sectional ovarian cancer data

We next apply HyperTraPS to a well-studied dataset on copy number variation in ovarian cancer progression (Knutsen et al., 2005). This dataset is included in the Oncotrees package (Szabo and Boucher, 2002), with (Loohuis et al., 2014) and we make direct comparisons these approaches. The data consist of a sample of *N* = 87 patients with differing levels of copy numbers for *L* = 7 genes associated with ovarian cancer, with the assumption that none of the alterations were present in the individual at birth.

FIG. 3A provides a visual representation of the dataset, showing the presence/absence of each genetic alteration in each patient. Fig. 3B shows the recorded transitions following parameter inference on the hypercube. A collection of previously unreported dynamic features are revealed by this approach. A set of several constrained, well-defined paths are visible, with flexible ordering in the acquisition of features being apparent. Interestingly, the feature that is acquired first has substantial influence over the subsequent pathway structure, visible as the tightly constrained individual pathways in Fig. 3B with rather few transitions between pathways. 3C shows the inferred ordering of trait acquisition in this dataset. Clear differences in the temporal order where features are acquired are observed here too. In both plots, alterations *8q+* and *3q+* are often observed early (the dark edges in the top left part of Fig. 3A, while *8p-, Xp-* and *1q+* tend to be acquired later. These observations are supported by Desper et al. (1999) and Loohuis et al. (2014). It should be noted that there is weak multimodality exhibited in almost all the features, consistent with the appearance of several distinct pathways in Fig. 3B.

**Fig. 3:**
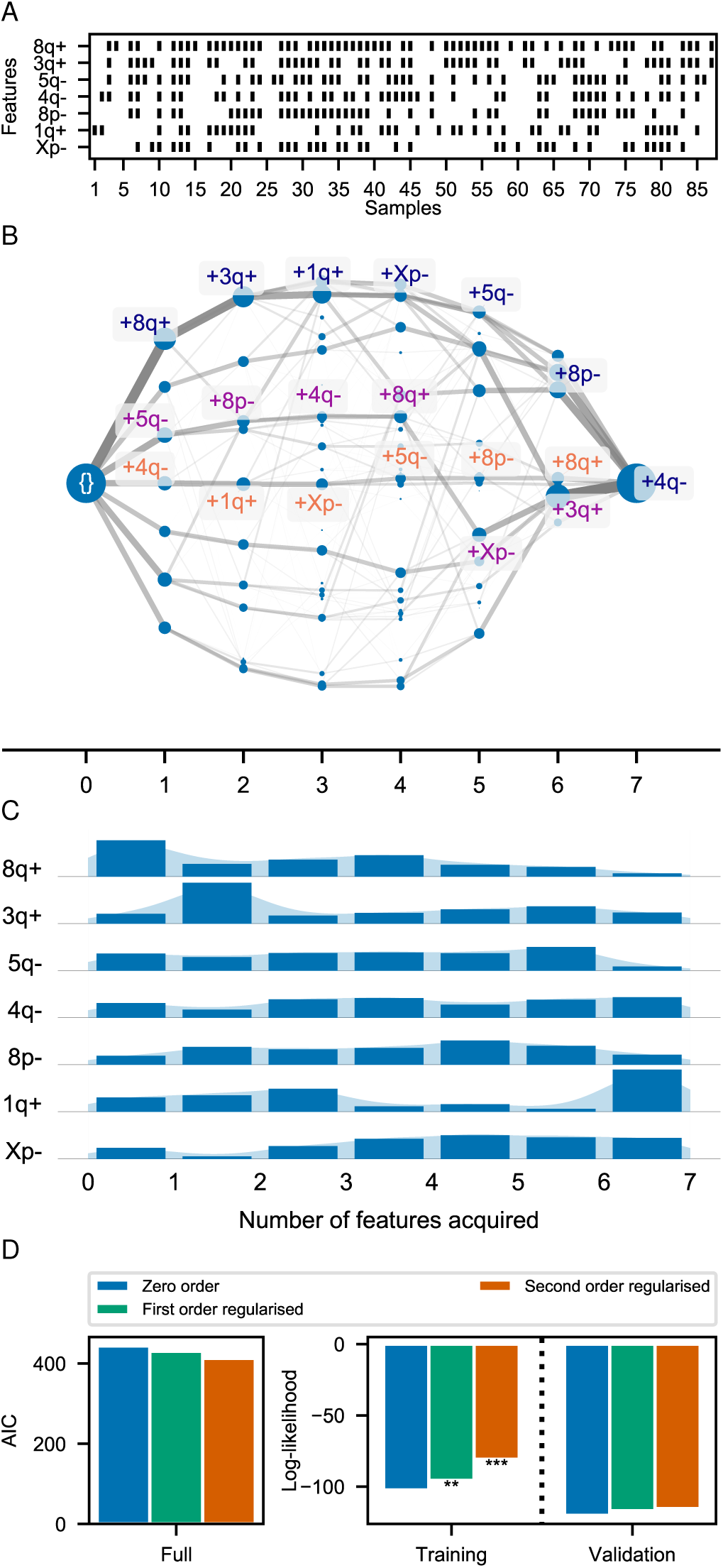
Inference, regularisation, and validation of HyperTraPS applied to an ovarian cancer alteration cross sectional dataset. A. The cross-sectional samples from the oncogenetic dataset are shown with black rectangles illustrating the presence of a genetic alteration in each sample. B. Trajectories across the hypercube from a healthy initial state to increasing number of copy number variants. Individual pathways are labelled with their cumulative steps: for example, the path {}→+8*q*+→+3*q*+ begins at the (empty) initial state, then proceeds first to the 8*q*+ state, then to 8*q*+, 3*q*+. Areas of nodes and edge weights are in proportion to the proportion of random walks through each. C. The inferred ordering histogram for the acquisition of genetic alterations. Blue bars give the probability that a given mutation (horizontal axis) is acquired at a given step (vertical axis). D. AIC for the full dataset and log-likelihoods for the training and validation data. The AIC score for the second order regularised model has the marginally lowest AIC, suggesting the importance of interaction terms. This is highlighted by the most significant training fit and the largest likelihood for the validation data.

In Fig. D.12A(iv) and B(iv), we compare the Capri results for the ovarian cancer data with the pathways inferred by HyperTraPS. Capri yields a DAG of causalities that matches the ordering of the most pronounced modes (and some of the most probably specific pathway structures) in the HyperTraPS posterior. HyperTraPS additionally reports alternative pathways supported by the data, which may be neglected in the maximum-likelihood Capri approach due to their comparatively low probability. The separation of these pathways (the aforementioned dependence on the first-acquired trait) is revealed through the HyperTraPS approach. For example, HyperTraPS suggests that while the *8q+* change is the most likely to be acquired first, if an alternative is acquired first, *8q+* can often be acquired rather later in disease progression. We consider additional comparisons with a Bayesian network approach in Appendix D.

In Fig. 3D, the AIC for each model is plotted along with the log-likelihood for the training and validation datasets. The second order regularised model outperforms the others with the validation dataset in terms of log-likelihood and significance. Additionally, the second order regularised model has marginal improved predictive power over the first order regularised and zero order (null) models with the validation dataset.

These results together illustrate that samples parameterisations from HyperTraPS model does provide improved predictive potential for a cross-sectional dataset with biomedical application.

### 2.3 Application to the evolution of multi-drug resistant tuberculosis

In the above two sections, observations are cross-sectional and ‘historically’ independent: each observation corresponded to an individual with its own unique trajectory from the ‘no features’ state on the hypercube. As discussed in the introduction, phylogenetically linked observations raise the possibility that traits may be inherited from ancestors rather than being acquired *de novo* for individual observations. Next, we demonstrate the application of HyperTraPS to a large-scale dataset with such a phylogenetic relationship between observations.

We consider the case of pathways of genetic polymorphisms that underpin drug-resistant tuberculosis isolates reported in Casali et al. (2014). In this study, the authors sequenced 1000 drug-resistant tuberculosis isolates from Samara in Russia. The data consists of presence/absence markers of polymorphisms at 16 key genes/promoter regions that confer drug-resistance, as well as mutations in three RNA polymerase genes for 993 of these isolates (those with complete data out of a total of 1000). Observed isolates are linked by a phylogeny, which Casali *et al*. constructed from genome-wide information (importantly, consisting of a much wider set of genomic regions than just those involved in drug resistance). We show the structure of this dataset with phylogenetic linkage (the maximum likelihood phylogeny from Casali et al. (2014)) in Fig. 4A.

**Fig. 4:**
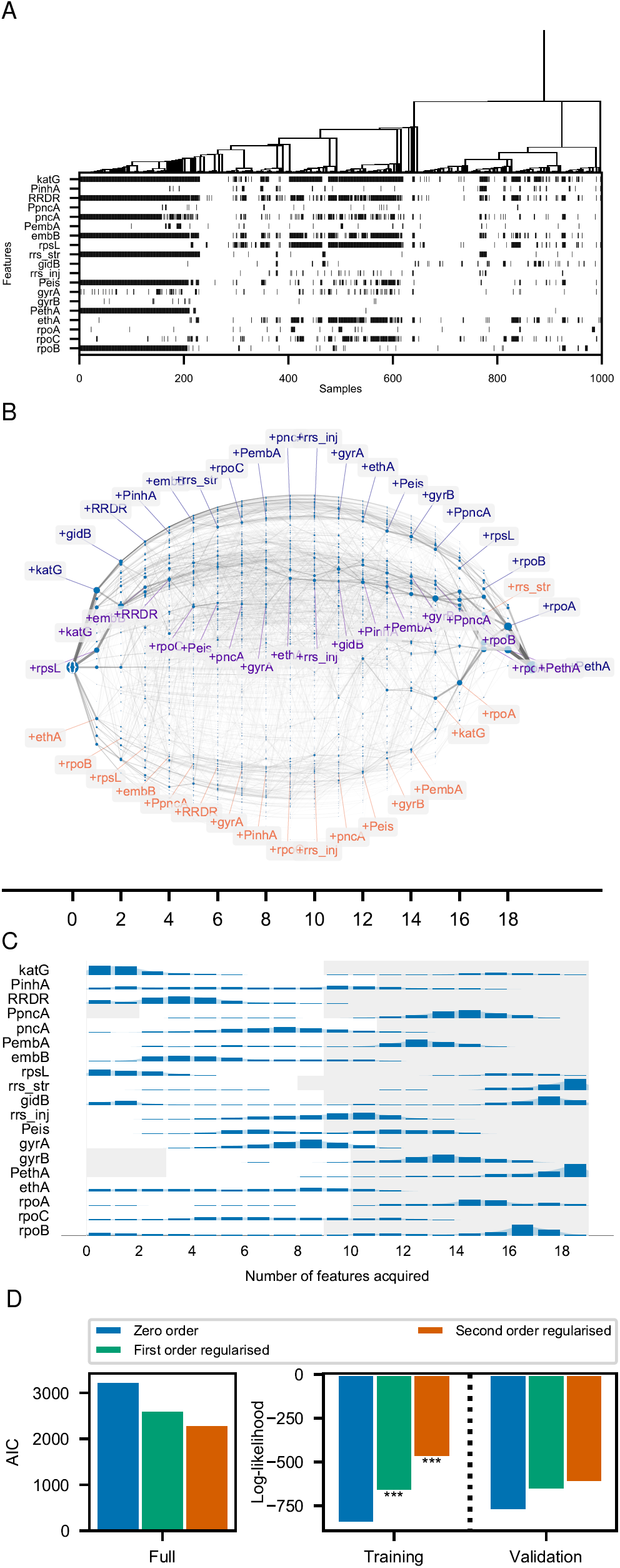
Inference and validation of HyperTraPS applied to genetic polymorphisms in drug resistant tuberculosis isolates. A. Each tuberculosis isolate (sample) is shown with a polymorphism at each genetic site indicated by a black rectangle in each column, linked by a phylogeny. B. Trajectories across the hypercube as a weighted directed graph using transitions between states simulated with posterior samples. There are clear differences in progression highlighted by the different regions of dense transition edges, with the first, second and sixth greedy paths labelled. C. Inferred ordering of changes to genes conferring drug resistance in tuberculosis. Bimodal ordering distributions reveal distinct evolutionary pathways underlying the acquisition of drug resistance. Density in the grey regions corresponds to acquisitions that do not directly affect the likelihood, as features are not observed to be acquired in these regions in the dataset. D. Regularisation of the model validation. The second order regularised model has the lowest AIC for the full dataset. The log-likelihoods for each model with the training and validations datasets are shown with all models experiencing statistically significant support over the null. The second order regularised model performs best for the validation dataset.

In addition to irreversible acquisition, we here assume that mutations are sufficiently rare such that convergent evolution is not a leading-order dynamic process between descendant and parent nodes in the phylogeny. With these assumptions, one can work backwards through the phylogeny and apply the bitwise AND operator between descendant states to allow a parsimonious estimate of the unobserved parent state. From these estimates, we can reconstruct the observations as the transitions from parent nodes to descendant nodes on the phylogeny which provide an independent set. A simple version of this process is depicted in Fig. 1A(iii). Once *D*^transitions^ has been reconstructed via the above process, the derived dataset can be used for inference of evolutionary pathways for the drug-resistant sites.

FIG. 4B shows a transition graph of the transitions across the hypercube with parameterisations from the inferred posterior. Once more, a collection of previously unreported dynamic features are immediately observed. There is substantial heterogeneity in density of edges in the transition graph and acquisition orderings between the different traits we consider. In contrast to the large number of highly focussed paths inferred from the ovarian cancer data, this transition graph demonstrates a smaller number of looser – but still distinct in structure – paths across the hypercube, each with a ‘cloud’ of variability indicating some flexibility in specific orderings within these pathways.

FIG. 4C shows posterior distributions on acquisition ordering, some of which, notably, are observed to be bimodal in time. This bimodality points to the presence of multiple pathways towards drug-resistance, with some traits having the potential to be acquired early or later in the evolutionary pathway. Each of these distinct modes suggests the presence of a distinct evolutionary pathway, supporting a picture where, for example, *katG* alterations may be acquired either early or late in resistance evolution. This presence of multiple distinct evolutionary pathways suggests substantial flexibility in the evolution of resistance (supported by the recent review of drug resistant tuberculosis in Dookie et al. (2018)).

The non-grey regions for each row of the histogram displayed in Fig. 4C represents the range over which each feature is observed to be acquired in the transition dataset, while the grey regions are the converse where acquisitions are outside the window of where the feature is observed to have been acquired in the data. As a result, density in the grey region is not directly supported by the data. However, under the assumption of full acquisition of all features provides a relative indication of when this acquisition would occur. This is further discussed in Appendix B where an alternative use of the posterior inference is presented.

Regularisation of the model finds that the minimum AIC arises at *k*_min_*≈*216 parameters out of the *L*^2^ = 361 starting parameters. Many interaction terms are thus required in the second order model, to provide significant explanatory power to distinguish multiple progression pathways (and the necessitated interactions between traits, as discussed above). Correspondingly, the AIC and log-likelihoods associated with the second order model are much larger than for the first order model. This observation aligns with the observation of bimodality in the ordering posteriors: interaction terms are required to enforce distinction between different progressions.

Validation calculations for the model (Fig. 4D) further support this message. All models experience statistically significant support over the null model in terms of the log-likelihood ratio. While the first order regularised model has improved predictive power over the null model, the second order regularised model provides around twice the increase in log-likelihood compared with the first order model.

In summary, the inferences around the order in which features are acquired from combining the phylogenetic relationships of samples yields new insights into the structure and variability of the evolutionary trajectories by which drug resistance is acquired. This can potentially add important insight into the mechanisms by which co-associating drug-resistant polymorphisms occur in an epistatic fitness landscape. We discuss some of the specific evolutionary implications in Appendix E and compare with outputs of the approach of Bollback (2006) in Appendix F (see Fig. F.14).

## 3. Discussion

In this work, we have introduced a powerful and highly generalisable statistical platform for inferring stochastic, coupled dynamics from samples in a binary state space. The methodology makes use of a stochastic process model on a hypercube, with Bayesian inference utilised to infer parameters from the full posterior with the HyperTraPS algorithm. We provide novel means for visualising the inferred paths across the hypercube, and introduce additional methods for working with inferred parameterisations including a framework for regularisation and validation predictions for choosing from a hierarchy of models. We have illustrated the method through application to synthetic datasets, before showing its utility and generality in application to real datasets: firstly, with oncogenetic data to construct a stochastic cancer progression model for the accumulation of alterations in an ovarian cancer dataset and secondly, with genetic mutations in drug-resistant tuberculosis to elucidate the order in which mutations are occurred that confer such drug-resistance. In each of these cases, new insight into the structure and variability of progression pathways was revealed through the inference platform.

The HyperTraPS platform is specifically designed to infer dynamic pathways given arbitrarily linked observations of many, possibly interacting, traits. The generality of this question is illustrated by the diversity of existing approaches that have some bearing on the corresponding inference problem. Table 1 illustrates several broad classes of these approaches, including regression models, Bayesian network models, stochastic processes on phylogenies, topological approaches and finite state space models (HyperTraPS).

**Table 1:**
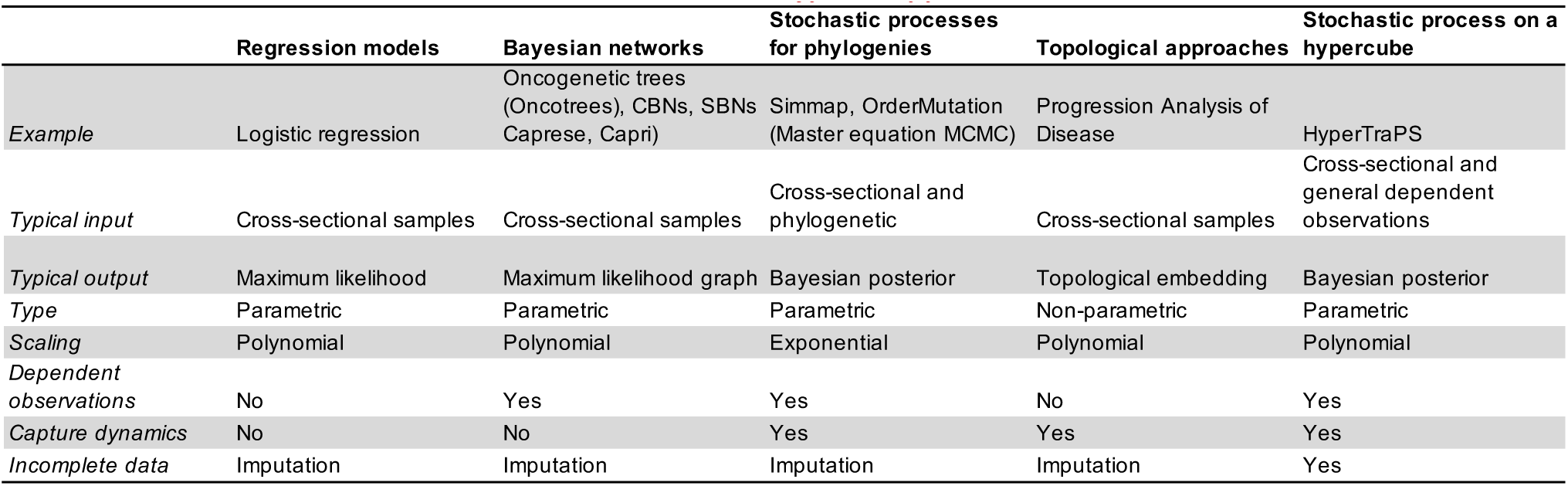
Comparison of HyperTraPS with other methods for inference from state space observations. We consider some of the key properties that HyperTraPS introduces. The following abbreviations are used: Suppes-Bayes Network (SBN), Conjunctive Bayes Network (CBN) and Markov chain Monte Carlo (MCMC).

| Types of approach |  |  |  |  |  |
| --- | --- | --- | --- | --- | --- |
|  | Regression models | Bayesian networks | Stochastic processes for phylogenies | Topological approaches | Stochastic process on a hypercube |
| *Example* | Logistic regression | Oncogenetic trees (Oncotrees), CBNs, SBNs (Caprese, Capri) | Simmap, OrderMutation (Master equation MCMC) | Progression Analysis of Disease | HyperTraPS |
| *Typical input* | Cross-sectional samples | Cross-sectional samples | Cross-sectional and phylogenetic | Cross-sectional samples | Cross-sectional and general dependent observations |
| *Typical output* | Maximum likelihood | Maximum likelihood graph | Bayesian posterior | Topological embedding | Bayesian posterior |
| *Type* | Parametric | Parametric | Parametric | Non-parametric | Parametric |
| *Scaling* | Polynomial | Polynomial | Exponential | Polynomial | Polynomial |
| *Dependent observations* | No | Yes | Yes | No | Yes |
| *Capture dynamics* | No | No | Yes | Yes | Yes |
| *Incomplete data* | Imputation | Imputation | Imputation | Imputation | Yes |

Regression models are widely used to perform classification and predictions, but the inclusion and/or reconstruction of dynamic observations is less explored in this framework. While this approach is applied widely across the statistical and biomedical community, it is usually reliant on a linear underlying model and does not attempt to capture dynamics in which variables arise. Additionally, it requires a clear dichotomy between predictors and response variables to be imposed *a priori*, when such a distinction may not be appropriate, especially from the perspective of the inference of pathways.

Bayesian networks allow the probabilistic relationships between features to be considered. They provide a common platform for this with different assumptions placed on the conditioning probabilities: two examples being Conjunctive Bayes Networks (Beerenwinkel et al., 2007) and Suppes-Bayes networks (Loohuis et al., 2014). These are commonly used in oncogenetic inference problems, and have proved successful at unpicking causal relationships between features. We have shown that HyperTraPS aligns with the outputs of these approaches in simple cases. In more general settings, the stochastic model underlying HyperTraPS has the potential to reveal more probabilistic structure, including the identification of competing stochastic pathways, complex sets of interactions between coupled traits, and the quantification of uncertainty in the pathway structures that are revealed.

Modelling trait evolution on phylogenies is the closest group of models to which HyperTraPS is related, and typically requires computation of master equation rate matrices that do not place restrictions on the transitions that may occur in the state space (Bollback, 2006; O’Meara, 2012). By embedding transitions on a hypercubic graph, HyperTraPS has the ability to handle orders of magnitude more features without noticeable loss of generality (simultaneous transitions are represented as equally weighted, temporally adjacent, transitions). Additionally, these methods are designed specifically for phylogenies, while HyperTraPS has applicability to generic sample dependency.

Dimensionality reduction approaches have been considered for finding representations of temporal dynamics from samples. Such methods are powerful and have been applied to vast data collected in whole genome single cell RNA experiments (Campbell and Yau, 2016) and also to disease (Nicolau et al., 2011). While highly flexible, these approaches often rely on specific assumptions about the quantitative details of the dimensionality reduction, leading to variability from method to method. Additionally, this approach is yet to be considered in detail for finite space state models like the ones we consider here.

The HyperTraPS framework presented here has several advantages: (a) its polynomial scaling allows it to deal with large (many observations and many traits) datasets; (b) the regularisation processes we outline allow it not only to reveal and deal with arbitrary coupling between traits, but to select good and statistically significant parametric representations of these couplings to yield sparse models (thus applying Occam’s razor); (c) it yields general and readily interpretable predictions; and (d) it simultaneously provides inferred pathway structure, mechanistic insight, and uncertainty quantification. A further advantage, which we have not explored in detail here, arises from its inherently Bayesian nature: the ability to include prior information about pathway structure. Throughout this article we have assumed uninformative uniform priors over pathway structure. In situations where, for example, the scientific literature provides existing insight into a pathway, the prior distributions on the hypercube parameters can readily be adapted to include this prior knowledge in the inference process.

Despite these advantages, there are of course some limitations to the platform’s capabilities. Incomplete data currently provides a challenge for inference with HyperTraPS. There is nothing in principle preventing *hypercubic inference* with incomplete data: unbiased random walks can be simulated on a hypercube and their ability to recapitulate observations can be computed. Indeed, HyperTraPS can be applied in the case of uncertain *end points* of observed transitions. However, the sampling algorithm that allows HyperTraPS’ efficient sampling of high-dimensional spaces currently does not translate to incompletely described *start points* of observed transitions. This is due to the complexity that arises from the potential compatible emission patterns with the source state producing differential weightings for the compatible transitions that contribute to the likelihood. The datasets considered here involved *complete* entries – we excluded cases where observations of particular feature in a sample were not present (this involved excluding 7 out of 1,000 samples for the case of tuberculosis). It is worth noting that the potential to handle incomplete data in the target state already is a significant advantage over many other methods where subjective imputation methods are required to make progress, and that incomplete data can be handled here for any case of independent trajectories where a complete source state is known, such as for cross-sectional datasets. However, further developments in this area would be invaluable for further generalisations.

We introduced a method for regularisation in HyperTraPS models through use of a greedy backward selection algorithm allowing the discovery of sparser model representations and to uncover the importance of interaction terms for a given dataset. This process both enables clearer insights to be drawn between the relationship of features for the progressions that best explain the data, while also providing a means of considering the extent to which the data is suggestive of multiple progressions. While this approach was successful towards the above, the means by which it achieves this was reliant on both the imperfect greedy algorithm and additionally on the subjective use of the Akaike Information Criterion (AIC) for finding such sparser models. A multitude of methods are available for performing model selection within a full Bayesian setting (O’Hara and Sillanpää, 2009; Murphy, 2012) and exploration of alternative approaches for exploration of mappings from *W→π* and regularisation of HyperTraPS models is an import avenue of research as has been the case across statistical modelling.

Our platform occupies the under-explored intersection between methods for inferring dynamics from uncoupled observations (as in cancer progression) and from phylogenetically linked observations (as in evolutionary inference). In addition, as we have demonstrated, HyperTraPS can harness both cross-sectional and longitudinal observation data. We have shown that HyperTraPS has a unique power to dissect multiple competitive dynamic pathways (yielding new insight in two biomedical case studies), and demonstrated how the processes of regularisation can be used to identify model structures that contain necessary and sufficient information to describe the set of pathways likely to exist in a given scientific setting. We anticipate that this flexibility, and the abilities of HyperTraPS to naturally quantify uncertainty and form probabilistic predictions about future behaviours, will be of use across biomedical and other scientific disciplines as volumes of data continue to increase.

## 4. HyperTraPS package implementation

All computational work was performed with custom-written software in C++ and Python. The code for the HyperTraPS package is available from the authors upon request.

## 5. Acknowledgements

All authors acknowledge support from EPSRC grant EP/N014529/1. IGJ acknowledges support from the University of Birmingham via a Birmingham Fellowship and from the Alan Turing Institute.

## 6. STAR Methods

### 6.1 Overview of HyperTraPS pipeline

In Fig. 5, we provide a diagrammatic overview of the HyperTraPS pipeline. The different elements are described below.

**Fig. 5:**
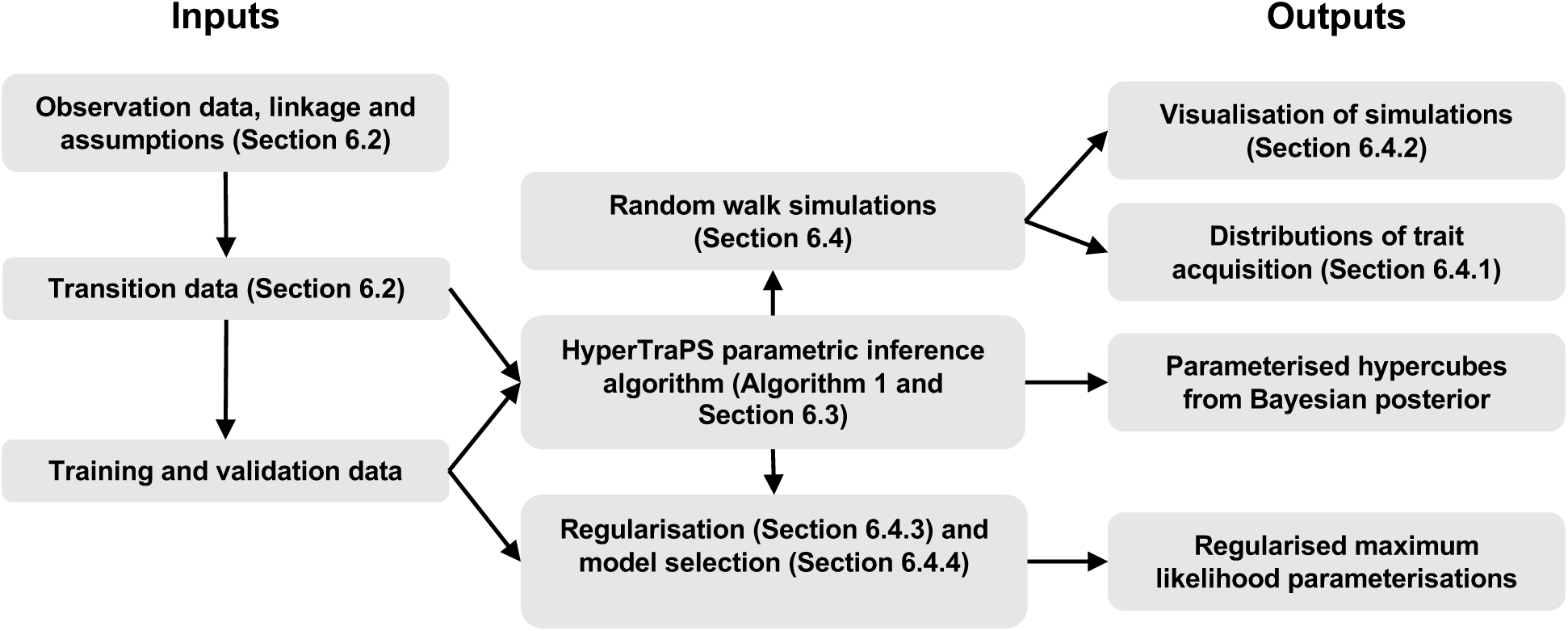
An illustration of the pipeline from inputs to outputs with the underlying inference, application and description methods within HyperTraPS.

### 6.2 Construction of a transition dataset from observed signals

As described in Fig. 1, the first step is to convert cross-sectional, longitudinal, or phylogenetically linked observations to a set of transitions, which we will represent as *D* = {*s*_*i*_, *t*_*i*_}, where *s*_*i*_ is the *i*th source state and *t*_*i*_ the *i*th target state, and there are *n*_*D*_ observations in total.

### 6.3 Inference of parameterisations

#### 6.3.1 Bayesian framework and likelihood of transition dataset

As introduced in Johnston and Williams (2016), we choose a Bayesian framework for inferring parameters for the set of edge weights *W* on the hypercubic transition graph that explain the data *D*. As such we are concerned with drawing samples from the posterior:

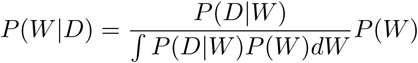

which is proportional to the product of our prior probability density *P* (*W)* on edge parameterisations and the likelihood ℒ (*W|D*) = *P* (*D|W)*, such that we have *P* (*W|D*) ∝ℒ(*W|D*)*P* (*W)*. Throughout this work we choose a uniform prior distribution on *P* (*W)* and therefore only need to consider the calculation of (*W D*) in order to derive samples from the posterior probability distribution.

From this transition set, we can decompose the likelihood into the following form (regardless of whether the source data was cross-sectional, longitudinal, or phylogenetically coupled Johnston and Williams (2016)):

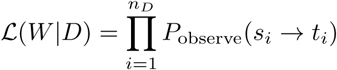

where *n*_*D*_ is the size of the transition dataset. *P*_observe_, the probability of observing such a transition requires a signal to be emitted by our system at both the source and target states, with the system having reached the source state and then made the transition to the target state via any possible walk on the hypercube. Therefore, the probability of observing such a transition can be written as:

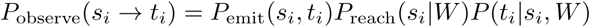

If each *s*_*i*_ and *t*_*i*_ is completely given (no missing data), emission probabilities corresponding to the observed transitions are independent of the trajectories taken, and *P*_emit_ yields a constant multiplicative factor which can be ignored in the inference process. In Johnston and Williams (2016), it is shown that the remaining log-likelihood can be written as:

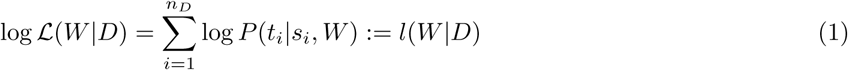

where the only computation required is the probability of making the transition to *t*_*i*_ from *s*_*i*_ for a given parameterisation of *W*.

In order to calculate *P* (*t*_*i*_*|s*_*i*_, *W)*, a sum over all possible paths between *s*_*i*_ and *t*_*i*_ is required. Given the number of paths between *s*_*i*_ and *t*_*i*_ scales as the factorial of the Hamming distance, the problem of deriving the rate matrix becomes intractable for systems of dimension *O*(10^1^). Instead we tackle the problem by way of performing biased random walks restricted to pathways that end in *t*_*i*_. This method of sampling was introduced in Johnston and Williams (2016) and allows systems more features to be considered than previously has been the case. This HyperTraPS algorithm that forms the key part of the HyperTraPS framework is captured in Algorithm 1.

**Algorithm 1:**
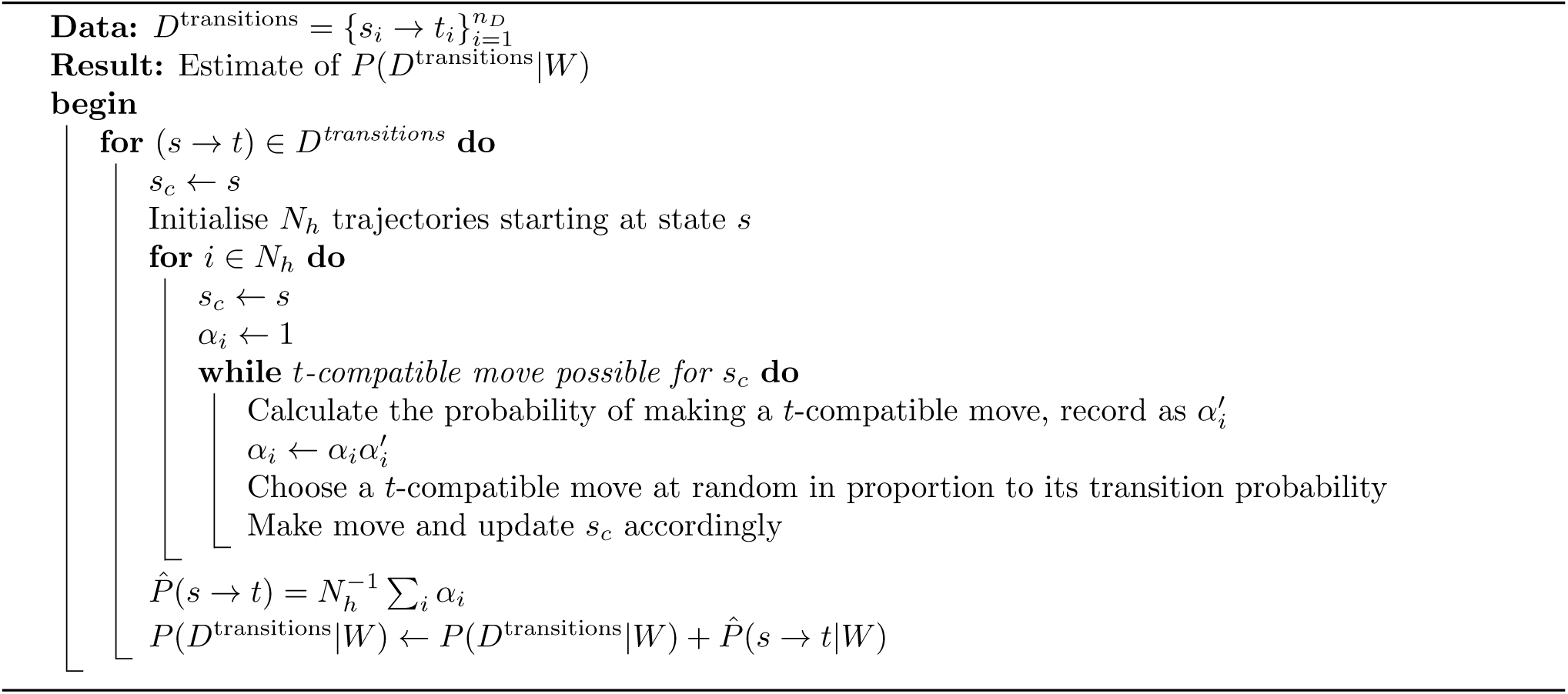
HyperTraPS algorithm for complete data: Hypercubic Transition Path Sampling was first introduced by Johnston and Williams (2016) to sample random walks on a hypercube across a restricted set of compatible states between a source and target state.

#### 6.3.2 Tractable parameterisations of hypercube

The hypercube embedding for *L* features has *L*2^*L-*1^ edges that we aim to parameterise. As *L* grows, we require a way of reducing this number of parameters *k* without compromising our ability to describe the dynamics of a system. Shrinkage and model selection tools may be used to achieve this reduction: we explore a simple approach for this process later. However, given the potentially large number of parameters in the default model, we also consider methods to reduce parameter space before the inference process.

One intuitive approach is based around considering the factors that may influence a given transition. The full parameterisation allows independent rates between any two states. In this picture, the probability *P* (*i*) of acquiring the *i*th trait can take arbitrary and independent values for every possible combination of the other *L–*1 traits. As an alternative, we can restrict the dependence of *P* (*i*) on the *coupling* of other trait patterns. For example, if we assume that each of the *L–*1 other traits influence *P* (*i*) independently (no synergistic interactions), we need only *L*^2^ parameters: a ‘basal rate’ of acquisition for each trait *i*, and the amount by which this basal rate is modified by the presence of trait *j* ≠ *i*. This reduction is analogous, for example, to Generalised Linear Models where response variables can be considered a function of independent variables and interaction terms between the independent variables, neglecting higher order interaction terms.

From this perspective a hierarchy of models may be constructed (Fig. 6). For the ‘zero order’ model every feature has equal probability of acquisition (*k* = 0 parameters). In the ‘first order’ model, every feature has an independent acquisition probability (*k* = *L* parameters). In the ‘second order’ model, every feature’s acquisition probability depends independently on the presence of each other feature (*k* = *L*^2^ parameters). Higher order models, including the full *L*2^*L-*1^ set can be envisaged, introducing more complex interactions between the co-occurrence of features.

**Fig. 6:**
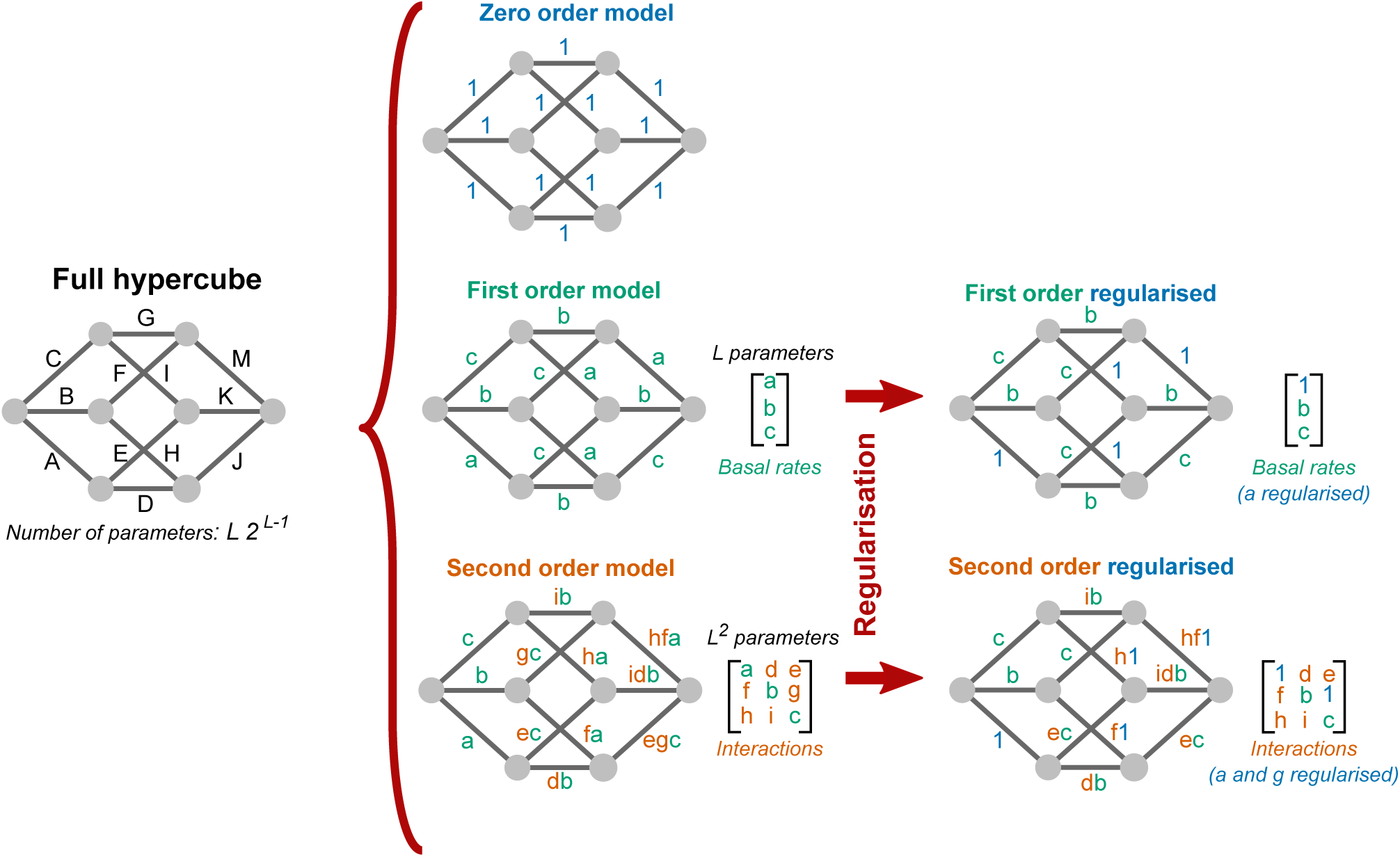
Tractable parameterisations and regularisation. A full irreversible directed hypercube is parameterised by edge set *W* and contains *L*2^*L-*1^ edges. We define three orders of model (*zero order, first order* and *second order*) for reducing the parameter space and regularised models (*first order regularised and second order regularised*). The zero-, first- and second-order models are nested in the sense that a second order model can be equal to a first order model (interaction terms all set to unity) and the first order model can equal to the zero order model (all basal terms set to unity). In the example above, for *L* = 3, the 12 edges of the Full (hyper)cube (A-K) are reduced down to combinations of a set of 9 parameters (a-i). The advantage becomes clear for larger *L*. At *L* = 16, over 500,000 potential parameters are reduced to just 256 parameters for the second order model. Regularisation involves a greedy backward selection process to identify which parameters may be removed (set to the value of the zero order model, unity) and decrease a criterion, which we choose to be the Akaike Information Criterion. In the illustration above, for the first order regularised model, parameter *a* is set to unity and, for the second order regularised model, parameters *a* and *g* are both set to unity (as would be the case in a zero order model) with the consequent impact on the hypercube edge weights shown.

A mapping is then required for applying the parameters in each model to the edges of the hypercube. We choose the following functional form:

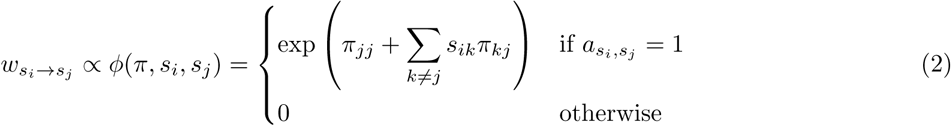

where 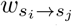 is the transition weight associated with going from state *s*_*i*_ to *s*_*j*_, *s*_*ik*_ is the *k*^*th*^ element of the state *s*_*i*_ and 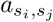 is the directed adjacency matrix of the hypercube. The choice of the exponential function in the mapping simultaneously enforces that *w* elements are non-negative and allows a computationally efficient coverage of a wide range of possible parameter values.

With the tractable parameterisation described, we replace *W* in the above inference with *π*, and consider the inference of the matrix representing the a priori model choice *π* instead of the full set of edges *W*.

At this point, it should be noted that this mapping retains the Markovian nature of walks on the hypercube as *W* is a static object independent of the walker. Alternative mappings can be construed that make the mapping a function of *the states previously visited* by a specific walker and update *W* for each walker. This introduces the idea of including non-Markovian dynamics within the same framework, with a similar hierarchy of potentially tractable parameterisations imaginable from this alternative perspective. However, this is beyond the scope of the work we introduce here.

#### 6.3.3 Monte Carlo sampling methods

The complexity of the inference problem challenges analytic or uniform sampling approaches to compute Eq. (1) for the full range of parameters *W*. Instead, we employ Markov Chain Monte Carlo (MCMC) in order to generate samples from the posterior on *W*. As the HyperTraPS algorithm generates an estimate of the likelihood (with the same expected value as the exact likelihood), this is in fact a pseudo-marginal MCMC sampler which has been shown to yield the same stationarity properties as if it were exact (Andrieu and Roberts, 2009).

Previous approaches for specific scientific questions (Williams et al., 2013; Johnston and Williams, 2016) found this pseudo-marginal MCMC sampler to demonstrate good mixing. However, there are cases where this simple approach produces poor mixing, specifically when the hamming distance between a source and target state becomes large. This is because Algorithm 1 generates an estimate of the likelihood with increased variance around its exact value due to the greater number of acquisitions made during path sampling. This can lead to poor mixing of sampler chains due to parameterisations becoming stuck in the tail of the estimator rather than being able to explore the variation of the likelihood with respect to different parameterisations. This occurs when the variance of the total log-likelihood has a variance with magnitude greater than unity (Sherlock et al., 2015).

To address this issue and generalise to more diverse datasets, we embedded HyperTraPS within an auxiliary pseudo-marginal MCMC algorithm (APM MCMC), which also satisfies the same convergence properties as MCMC (Murray and Graham, 2015). By making the likelihood a joint density *l*(*π, u*) over the parameters of the model and also the random variable from which our estimate is drawn, alternate Metropolis-Hastings steps can be performed by keeping *π* and *u* alternately fixed during the proposed update to the chain. For HyperTraPS, a new proposal for the random variable *u* is a new set of random trajectories across the hypercube over which each observations’s likelihood is estimated. We make use of this scheme throughout this work as little computational overhead is introduced with observed potential ability for improved mixing times.

As discussed, we have assumed a uniform prior for the parameterisations of the hypercube. For our choice of mapping *π*, this means we choose *P* (*π*) *∈U* (–*m, m*) where *m* = 10 covering 20 orders of magnitude with respect to the relative size of inferred parameterisations.

Rather than drawing from the prior distribution on *π*, for every Monte Carlo sampling run, we choose to begin from *π* = **0**. This is equivalent to the zero order model where there is no directionality pre-supposed. This facilitates the avoidance of local traps in the parameter landscape while remaining agnostic in introducing directionality into the inferred parameterisations for a particular dataset.

### 6.4 Applications of parameterisations

#### 6.4.1 Simulated walks to illustrate order of acquisition

The inference process above yields inferred posterior distributions on the hypercubic edge weights *W*. We can query these posteriors in a number of ways to gain descriptive and predictive information about the mechanisms generating observed states. First, we produce a parsimonious and intuitive representation of the dynamic pathways supported by the inferred posteriors. Here, we simulate an ensemble of random walkers generating complete trajectories on hypercubes with sets of transition probabilities sampled from the inferred posterior. This ensemble reflects the likely dynamic pathways supported by the dynamic transition model after parameterisation. We simulate an ensemble of random walks in two ways: *Walk Simulation 1 (WS1)*, with walkers that run from completely {0} ^*L*^ to {1} ^*L*^ where a feature is acquired at every time step and *Walk Simulation 2 (WS2)*, that run across the transition dataset *D*^transitions^. In each case, we record every transition between states allowing the construction of a weighted directed graph of all states and transitions encountered. From this graph, the frequency *f*_*ij*_ with which feature *i* is gained at step *j*. In the main text we focus on graph and histogram depictions of WS1 and refer to Appendix B for further discussion and illustration of WS2.

#### 6.4.2 A graph embedding and visualisation for dynamic acquisition on the hypercube

With each simulated random walk, *L* transitions occur between states on the hypercube. Across a large sample of these states we encounter a set of states 𝒮 = {*s*_*i*_} and we can represent the number of transitions between any two states by a directed, weighted graph with adjacency matrix *a*_*ij*_.

In order to visualise this graph to reveal characteristic progressions across the hypercube resulting from the parameterisation, we use a custom embedding to project the high-dimensional graph into two dimensions. First, we project the hypercube on to the surface of a sphere, an intuitive choice given that the discrete hypercube is embedded on a continuous hypersphere. We optimise the projection by making the following choices:

- Every state is on the surface, so is given the same radial coordinate, *r* = 1.
- The number of features acquired in the dataset is a measure of the how far the state is along the progression from 0^*L*^ to 1^*L*^. Therefore, for every 𝒮 state, we find the sum of features present and assign a polar angle that produces an even distribution between 0 and *L* across the sine of the polar angle which lies on the interval 0 ≤ *θ* ≤ *π*.
- The azimuthal angle *ϕ* on the interval 0 ≤ *ϕ* ≤ *π* is assigned by considering the *mean angle of the states from all incoming edges*, in order to attempt maximise the potential spread of the most common distinct paths across the hemisphere. A final assumption involves choosing all states with a single acquisition (*L* states) to be uniformly spread on the cosine of the interval [0, π].

With the embedding, the plot of the adjacency matrix *a*_*ij*_ is augmented by choosing node sizes and edge widths in proportion to the number of times the state and the transition are respectively encountered by the ensemble of random walks. Three examples of plots generated from this embedding with parameterisations of the hypercube are shown in Fig. 2B(i)-(iii), illustrating the potential highlight different underlying progressions inferred with HyperTraPS.

In presenting the embedding, we adjust the graphical depiction to highlight the features of the graph in the following way:

- Vertex area is in proportion to the number of times the vertex is visited by WS1 simulated random walks.
- Edge widths and opacity are in proportion to the number of times the transition between states is made with a random walker under WS1.
- In the main text, all vertices are blue. It is also possible to colour encountered states in accordance with the whether depicted states *s∈D*^transitions^. In Figs. C.9B, C.10B and C.11 we examine examples where such considerations provide further insight.
- A greedy labelling scheme is employed to aid understanding how random walkers acquire features across the hypercube. As walkers start from the empty “{}” state (0^*L*^), we can consider the addition of single features as each edge is traversed. In the plots, we use a greedy mechanism for determining which edges to label. Starting from 0^*L*^, we take the most probable outgoing edge at each vertex encountered and label the feature acquired across that edge at the resulting vertex until the 1^*L*^ state is reached, giving is the first greedy path. The following *n* greedy paths make use of the same approach but disregard any previously labelled edges, taking the next most probable available. We use the approach to clearly identify the left-right and right-left paths in Fig. 2B(i)-(iii).
- Finally, an optional transform to remove vertex overlap may be applied to remove overlap of vertices with a given number of features, while retaining the relative area of each vertex that is determined by the number of times the vertex is encountered.

#### 6.4.3 Regularisation

We previously discussed approaches to reduce the parameter space of the HyperTraPS model while retaining dynamic information. We can also employ model reduction approaches to identify supported parameter structures given a particular dataset. This regularisation helps identify more interpretable, parsimonious models and to guard against over-fitting.

One approach to model selection would be a fully Bayesian exploration of the joint space of model structures and parameters. However, the combinatorial explosion of search space with *L* currently makes this approach unfeasible for all but the simplest systems. Instead, we sacrifice a full exploration of this complicated space in favour of a tractable but principled approach to balance the reduction of model complexity against the ability to fit the data. This illustrative metric can indicate the amount of redundancy present in the parameterised *π* that can be removed in order to reduce the potential for over-fitting. To this end, we introduce a cost function to penalise the log-likelihood and then perform a algorithmic search to optimise this function.

We note that the number of parameters *k* required to adequately describe a given dynamic system is deeply related to the mechanisms underlying that system. If features are acquired independently, the first order model with *L* parameters should be sufficient to capture the dynamics. If a higher order model with more parameters is required, it suggests that interactions exist between features, such that one feature may influence the acquisition propensity of another. Identifying the sparsest model that can account for observations therefore also reveals mechanistic insight into the system.

For simplicity, we use the Akaike Information Criterion (AIC) (Murphy, 2012) to introduce sparsity. The AIC score for a model can be written as:

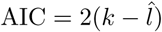

where *k* are the number of parameters in the model, and 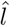 is the maximum log-likelihood. The score comprises the log likelihood and a penalty for lack of sparsity, in this case, the number of non-zero elements included in the maximum likelihood parameterisation *π*. Other options for regularization scoring include the Bayesian Information Criterion (BIC), but we refrain from exploring different metrics here, focussing firstly on illustrating how such regularisation can be performed within the HyperTraPS framework. A more general model selection approach will be the subject of future work.

To find parameterisations that optimise the AIC, we take a *greedy backward selection* approach (Murphy, 2012) to reduce the number of parameters *k* for a given model type. The process can be applied to both the first- and second-order models. An issue with such a greedy approach is that each single greedy backward step is unable to account for interactions between multiple parameters that lead to lower scores. Therefore, given a set of potentially distinct approximately maximum likelihood parameterisations, different backward selection processes from different starting maximum likelihood models may yield different minimum AIC scores for a given value of *k*. In an attempt, to bypass this problem, we take an ensemble of the top 100 maximum likelihood parameterisations from an MCMC sampling procedure and perform the greedy backward selection process to each one. Across the ensemble, for a given parameter number *k*, we take the minimum AIC score as a proxy for the minimum model at this level of parameterisation. The global minimum with respect to AIC is taken as the *first order regularised* or *second order regularised* model for the a first order and second order starting point respectively. The regularised models are then taken used in the subsequent section to perform model validation.

In Fig. 2D(i)-(iii), we show the regularisation process described above for the minimum of the ensemble at each value of *k* for the synthetic datasets and, in Appendix A, the process for ovarian and tuberculosis datasets. In these cases, a Hanning-tapered convolution is applied for smoothing to remove any artificial lack of the a single global minimum that should be expected.

#### 6.4.4 Validation

Importantly, the inferred parameterisations from our approach can be used to predict future behaviour of for a given state. We have described two procedures for generating parameterisations: sampling from the full posterior for a given model (first- or second-order) or regularised parameterisations constructed by the procedure in the previous section. In this section, we perform model validation through using the regularised parameterisations in order to identify the strength of evidence for the first- or second-order models. Using the outcome of this procedure, either samples from the full posteriors of the identified model or the corresponding regularised parameterisation can be used for prediction.

We validate this predictive power through two methods: firstly, through basic model comparison between the regularised first- and second-order models; and subsequently, by calculating the likelihood of observing data not used in the inference part of the method as a proxy for the predictive capability of each model. As a simple procedure to illustrate this, we split the *D*^transitions^ dataset into two halves: a training dataset *D*_train_ on which samples from the posterior are drawn and model comparisons can be made, and a testing dataset *D*_test_ with which the likelihood can be calculated using samples from the posterior for *D*_train_.

For model comparisons, we choose the zero order model as a null model. For comparisons between the different order models, we find the regularised first- and second-order model for the training dataset and denote this likelihood as 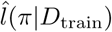. We then perform a log likelihood test, generating a log-likelihood ratio statistic (LLR) following a 𝒳^2^ distribution:

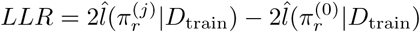

where 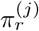 is regularised *j*^th^ order model. We compare to the 𝒳^2^ distribution for the number of non-zero parameters in *π*. With regard to the test dataset, we then use HyperTraPS to estimate 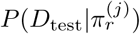 providing a measure of predictive capability of the *j*^th^ order regularised model, with larger values indicating better performance.

In each results section, a figure with the AIC score for each model, and the values from the model validation described above are shown, highlighting the performance of each model against the null model facilitating the determination of the appropriate model for use in a predictive setting for a given dataset.

## Appendix A. Regularisation of first order and second order models for ovarian and tuberculosis datasets

**Fig. A.7:**
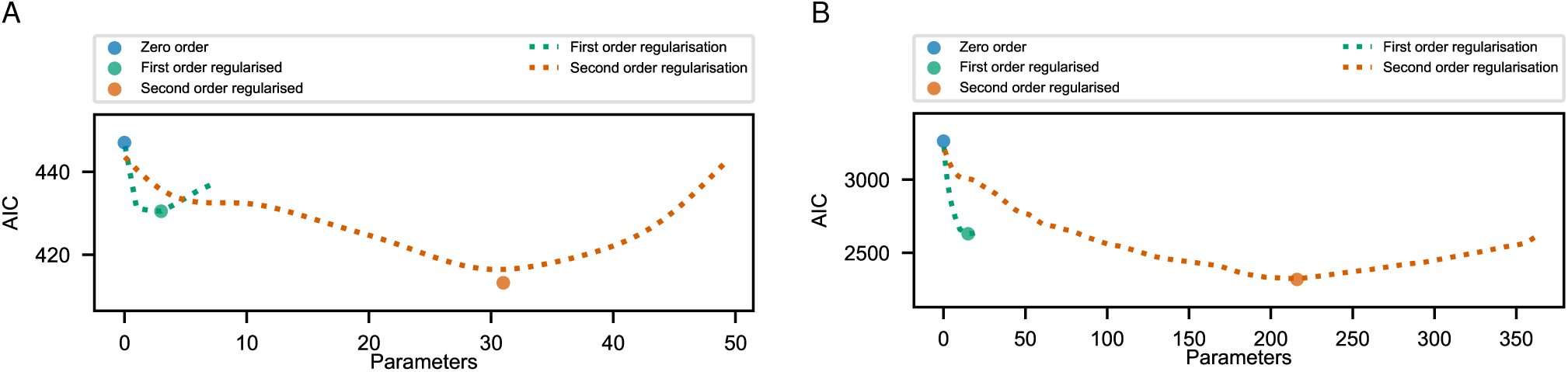
The regularisation process described in STAR methods and displayed for the synthetic datasets in Fig. 2 is applied to the ovarian and tuberculosis datasets. In both cases a smoothing algorithm (Hanning-tapered convolution) is applied to remove the appearance of local minima that would be unlikely to occur for larger ensembles of the greedy process. In both cases, the second order model is favoured, in particular for tuberculosis where a large number of the *L*^2^ parameters are retained with minima present at *k ∼* 216.

In STAR methods, we introduced a greedy backward selection process for inducing parsimonious parameterisations from samples of maximum likelihood models and demonstrated the process for an ensemble for the synthetic datasets (Fig. 2D). In Fig. A.7, plots for the ovarian and tuberculosis datasets are also shown with the minimum AIC score at each *k* from 100 unique greedy backward selection procedures for different maximum likelihood parameterisations. The AIC score is observed to decrease to a global minimum for each model. First order models may only have a few parameters removed before reaching a minimum, while second order models, depending on the number of interactions in the underlying dataset, can have a greater proportion of parameters removed. In both cases, smoothing is applied to remove the artificial appearance of more than a single minimum. This has no affect on the global minimum that is found from the ensemble and is used for illustrative purposes.

## Appendix B. Additional simulation for order acquisition interpretation: Walk Simulation 2

In Section 6.4.1 we introduced a protocol for using samples from the posterior of ℒ(*π|D*) to illustrate the order in which features are acquired. We denoted this process Walk Simulation 1 (WS1) as simulations from {0}*L* to {1}*L* are performed with the feature *i* acquired at step *j* being recorded as a proportion *f*_*ij*_. Alternatively, we can consider *f*_*ij*_ as the probability:

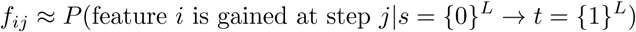

where *s* is the source state and *t* is the target state of the set of random walks.

As a feature is a always gained in each step, and all features are gained at some stage during this simulation process, the two properties ∑_*k*_ *f*_*kj*_ = 1 and ∑_*k*_ *f*_*ik*_ = 1 both hold. We illustrated the result of this simulation using a histogram for the matrix *f*_*ij*_ with kernel density estimates overlaid for each feature.

An alternative simulation protocol is to only simulate trajectories corresponding to transitions that are observed in the dataset. In other words, rather than assuming walkers proceed from 0^*L*^ to 1^*L*^, we simulate a set of walkers between each pair of source and target states *s*_*i*_, *t*_*i*_ in the dataset, relaxing the requirement that walkers start at 0^*L*^ and end at 1^*L*^. We denote this process Walk Simulation 2 (WS2). Here, we record the proportion of times that feature *i* is acquired at step *j* as *g*_*ij*_. However, in this case, *g*_*ij*_ does not satisfy the relations ∑_*k*_ *g*_*kj*_ = 1 and ∑_*k*_ *g*_*ik*_ = 1 anymore, as there is no explicit guarantee that feature *i* is acquired across all pairs of transitions {*s*→*t*}*∈D*^transitions^, nor that there is a transition at every step *j* observed across the transition dataset. Framing as a probability of acquisition of feature *i* at step *j* given it is acquired, we can consider WS2 as:

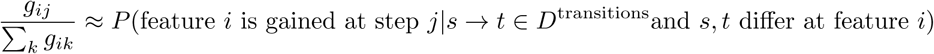

where *s* is the source state and *t* is the target state for each element of the set of simulated walks. This fraction is the proportion of times that feature *i* is gained at step *j* given it is acquired in a transition observed in the dataset.

The main distinction between WS1 and WS2 is the following: WS1 infers trajectories, informed by data, that start at {0} ^*L*^ and acquire all features to reach {1} ^*L*^. WS2 restricts the inference to the region ‘covered’ by the set of transitions observed in the dataset. Therefore, WS1 provides a readout of a complete process of acquisition (so may be more appropriate for analysis in systems where this is the expected outcome), while WS2 gives a readout of trajectories without extrapolating beyond the limits of observed states (and may be more appropriate if the walks are not believed to go to completion).

**Fig. B.8:**
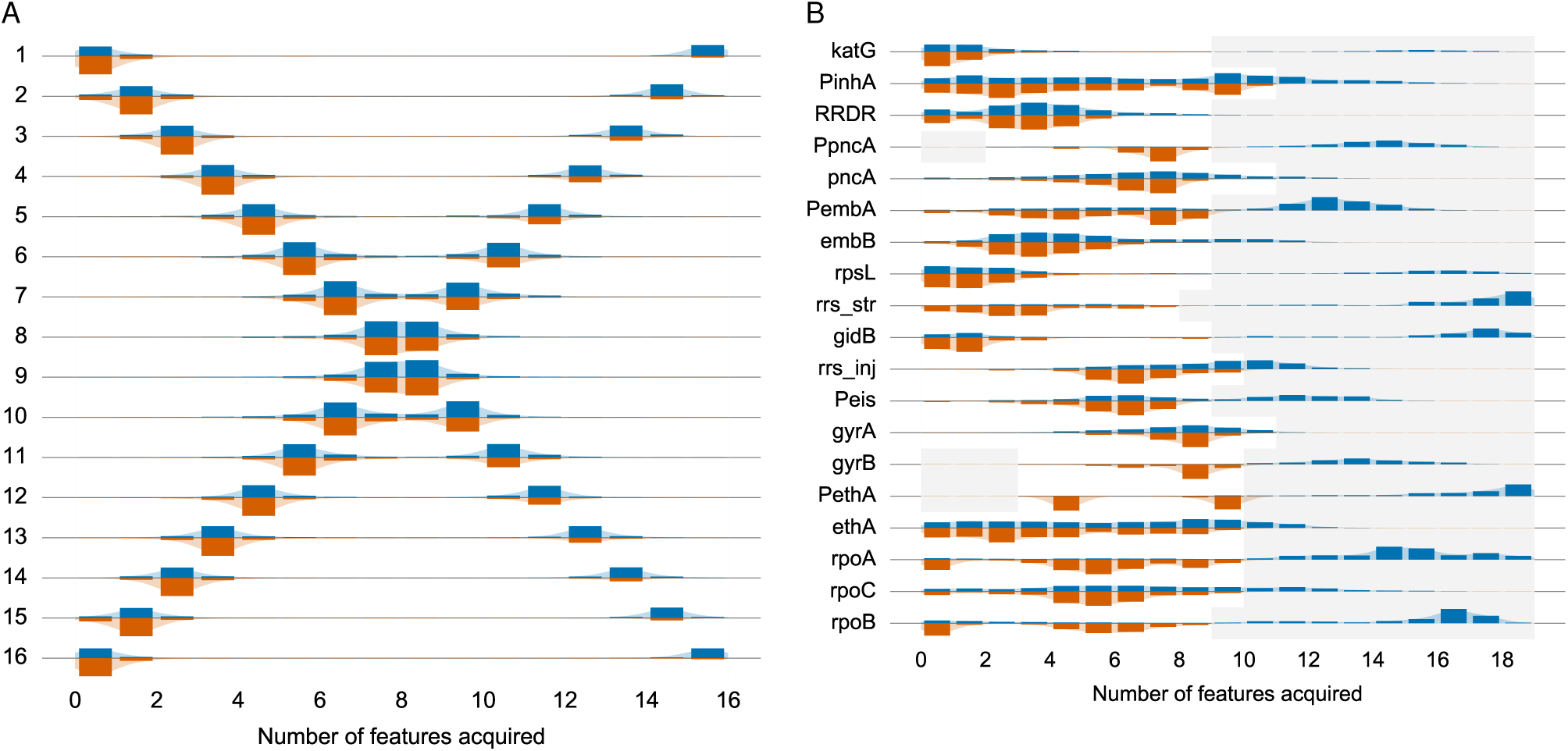
Comparison between WS1 and WS2 represented as violinplots for the cross-sectional dataset 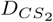 (A) and tuberculosis (B) from the main text. The blue bars are density corresponding to acquisitions with WS1 and the orange density for acquisitions with WS2. WS1 represents density for the ordering of acquisition from 0 to the 1 state so assumes all features are acquired. WS2 provides density relating to the acquisitions observed in the dataset.

We plot the densities for WS1 and WS2 in Fig. B.8 for two datasets from the main text, 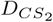 and the tuberculosis dataset. WS1 bars above the axis (blue bars) and WS2 bars below (orange bars) with kernel density estimates over each to guide the eye. As a result of this different approach, there are three key differences. First, posterior probabilities are rescaled according to how much a trait is ‘covered’ by observations. This is seen, for example, in feature 1 (and feature 16) in Fig. B.8A. Here, under WS1, early and late acquisitions of the feature are inferred to be equally likely, as walks are inferred to always run to completion. Under WS2, the number of walks that run to completion is lower (only some observations include ‘complete’ acquisition). The early acquisition mode is then inferred to be more likely, with a balancing probability that the feature is *not* acquired.

Secondly, with WS1, as the process starts from 0^*L*^, the transitions observed in the dataset are not guaranteed to be *reached* by random walkers. This means that the overall inferred parameterisations across the entire dataset may not lead transitions in the dataset being encountered. As a result, the WS1 process does not allow us to directly enquire into the nature of progressions between states in the original transition datasets. By exactly considering these transitions, WS2 allows this data to be examined using the parameterisations that have been sampled across the entire dataset allowing for a different type of inference. A clear example of this is seen in Fig. B.8B for feature *PembA* or *PethA* that are rarely encountered in the window of acquisition where they are acquired in the dataset, illustrated by the strikingly different distributions for WS1 and WS2.

Thirdly, there is no density observed in the grey regions for WS2 due to there being no transitions in the dataset ‘covering’ these regions, so no transitions performed with WS2 record any density there. In Fig. B.8B, in application to the tuberculosis dataset, the lack of WS2 density (orange bars) in the grey regions is apparent. In addition, there is clearly observable multimodality in WS2. Multimodality in WS1 is indicative of a feature belonging to multiple progressions that may include an absence of acquisition if the trajectory does not terminate. In contrast, multimodality in WS2 is indicative of multiple progressions where multiple orders of acquisition of a given feature are directly observed in the data. A striking example is *PethA* where in WS1 the predominant visible mode of acquisition is in the grey region towards the end of all possible acquisitions, while in WS2, the acquisition is observed in two distinct regions at step *j* = 5 and step *j* = 10, suggesting that the transition data contains multiple types of progression where *PethA* is acquired. This is also clearly the case for other features such as *PembA, PinhA, ethA* and *RRDR*.

We introduced WS2 here as a supplementary form of enquiry of the posteriors that can potentially reveal additional inferences about the underlying progressions from which the data may be derived. In the next section, we look in more detail at the assumptions, types of progressions and the outputs in the plots we have used for the inference in order to motivate intuition further.

## Appendix C. Implicit assumptions and interpretation of parameterisations

### Appendix C.1. Definitions and assumptions

In using HyperTraPS with a dataset of samples *D*^samples^ linked to form *D*^transitions^, several technical concepts and assumptions are required. We review these here:

- Systems are assumed to evolve as an ensemble of trajectories on a hypercubic transition graph. Independent sets of samples emerge from independent trajectories in this ensemble; historically coupled sets of observations (longitudinal or phylogenetic) observations emerge from trajectories that share some or all of their history.
- We assume that features are *irreversibly* acquired along a trajectory.
- As with WS2 above, progressions of a system may not always involve the complete acquisition of all features before an outcome occurs. For example, with respect to cancer alterations, not all mutations may be acquired before death occurs. In the tuberculosis example, not all polymorphisms may be required before drug-resistance is acquired. This assumption needs consideration when interpreting results as it may shape the research question being asked.
- *Observations* arise from signals emitted from states of the system where features have been acquired in a specific order.
- We assume that these signals faithfully reflect the state of the system when emitted. Below (‘Noisy observations’), we explore the effects of noise in these signals below and show that even in the presence of substantial observational noise we can recover the underlying generative pathway structure.
- We assume that the probability that a given state emits a signal is the same for all states. This assumption may be challenged, for example, if observations occur regularly in real time and some states are occupied for longer than others. However, we show below (‘Non-uniform sampling across the progression’) that HyperTraPS’ inference of event *ordering* (not explicit timescales, which we do not address) is robust to this class of challenge.
- The inference process yields a fundamentally probabilistic output: a posterior distribution describing the probability of different dynamic pathways. We interpret modes in this posterior as ‘characteristic’ pathway structures. Spread of posterior probability density around these modes corresponds to ‘variations on a theme’: pathways of similar structure with similar statistical support given the data. Below (‘Repeated uniform sampling’), we explore how the volume and structure of sampled observations affects this posterior breadth.

In the next section, we consider this set-up for the inference process by constructing specific synthetic crosssectional datasets to examine the impact of these assumptions on outputs.

### Appendix C.2. Implications of assumptions for HyperTraPS inference and walk simulations

To illustrate this further, we consider specific attributes with respect to attributes that may be present in systems under consideration:

1. *No structure*: only in the case of independent feature acquisition and identical frequencies will no suggestive progression be found, in which case the prior distribution (in this article, uniform across all trajectories) will be recovered by the inference process.
2. *Samples from complete and partial progressions*: If one or more of the underlying progressions does not correspond to a complete walk across the hypercube, transition density in unsampled regions will be dictated by extrapolated dynamics or the prior, depending on whether WS1 or WS2 is used. In Fig. C.9(i) we illustrate the dataset 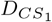 for *L* = 8 but for a progression that now stops after gaining feature *i* = 4. In this case, with no other progressions present in the dataset, we find that the remaining features gained in the grey region do so with a uniform distribution over remaining orderings (recovering the prior). In Fig. C.9(ii) we examine the case where there is a complete right-left path and a partial left-right path (that ends with feature *i* = 8 being acquired, which is the start of the complete trajectory). Trajectories belonging to the left-right transition in WS1 may be interpreted as joining the full right-left path. WS2 does not clearly disambiguate these dynamics – it is not clear whether features 5-8 are acquired. WS1, in the bottom right quadrant of the plot, shows some support for the beginning of the complete progression beginning after the partial progression ends. Fig. C.9(iii) looks at two partial progressions again illustrating that in the grey region (acquisitions without support in the dataset), there can be a mixed signal from the two partial progressions.

**Fig. C.9:**
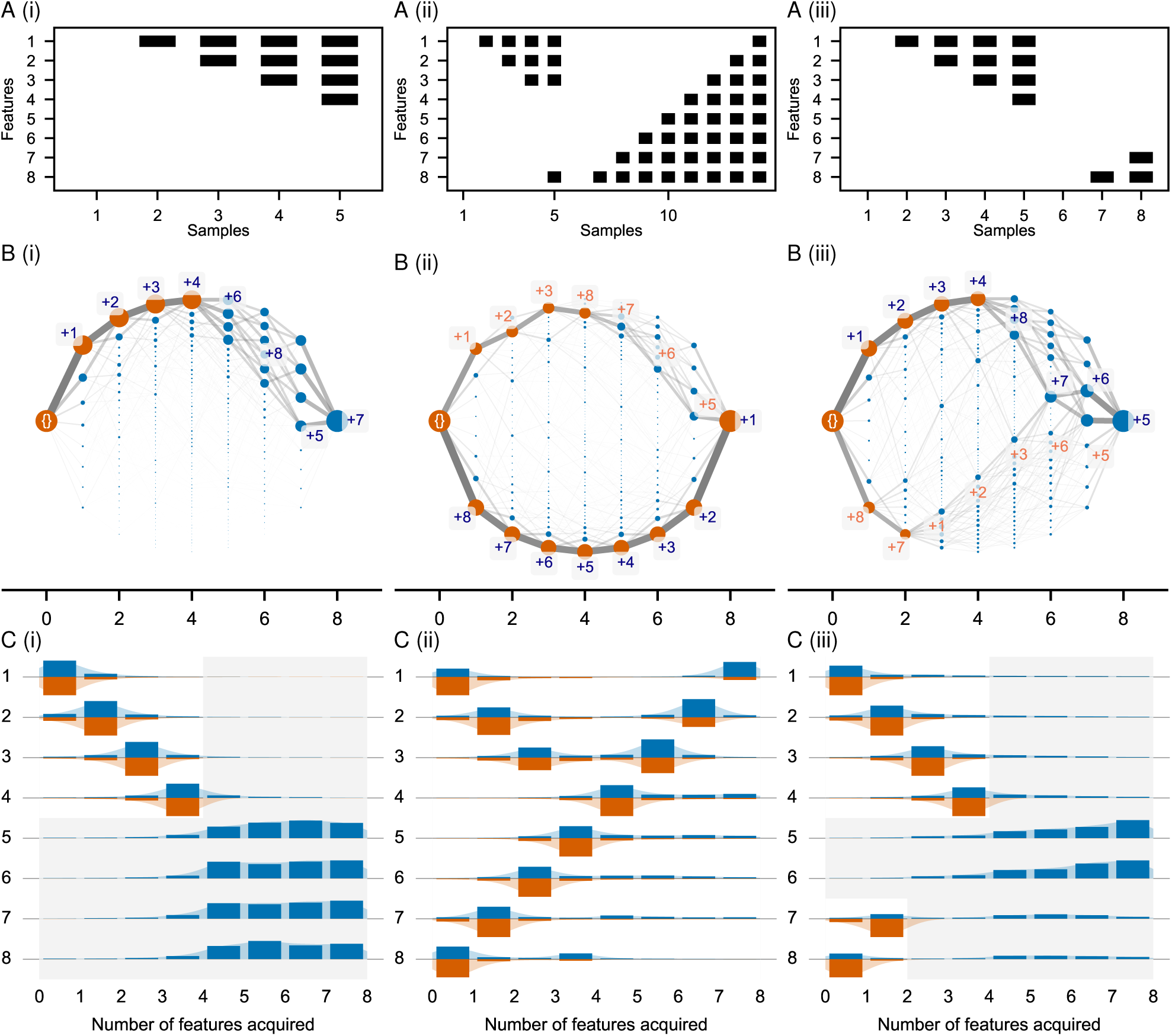
HyperTraPS inference in the presence of partial and multiple progressions. Three datasets are considered: (i) A single partial progression (1,2,3,4); (ii) A single partial progression (1,2,3,8) and a complete second progression (8,7,6,5,4,3,2,1); and (iii) Two partial progressions (1,2,3,4) and (8,7). In each case: A. shows the dataset structure (*dataset plots*); B. The inferred paths on the hypercube with samples from the second order posterior and WS1 simulations (*hypercube plots*). Orange vertices are observed in the dataset, while blue ones are not; and C. The corresponding histograms for WS1 and WS2 (*histogram plots*). For (i), the partial progression is inferred following by uniform acquisitions in line with the prior expectation. In the hypercube plots, paths on the hypercube are seen to diverge with equal proportion in this region illustrating this point. For (ii), the hypercube plot highlights the ability to infer both progressions. The longer path has greater weight due to an increased number of observations associated. The greedily labelled paths show an interesting feature where at the end of the partial progression, as the last feature is the first feature of the complete progression, the pattern of acquisition seen in the second progression is ‘predicted’ to occur in continued acquisition. This is visible in the histogram plot by the asymmetric density in WS1 flowing from feature *i* = 7 for the fifth feature acquired onwards. For (iii) with two partial progressions, the two paths are clearly distinguished in the hypercube plot with the same property of the progressions continuing on from each other after each partial progression is completed, eventually joining together after the sixth feature is acquired. The spread of other states encountered highlights the stochastic nature of the platform’s predictions.
3. *Noisy observations*: We consider the influence of noise in observations in Fig. C.10 by looking at the single left-right progression conflated with noisy observations (from a cross-sectional dataset made up of 10 randomly sampled trajectories). From Fig. C.10(i)-(iii), the number of noisy (random acquisition of traits) observations increases, introducing breadth into the inferred posterior around the modal pathway (Fig. C.10(iii) for example). However, even with 50% noisy observations in Fig. C.10(ii), it is possible to clearly recover the modal progression. Even for the extreme case, the non-noisy pathway is almost exactly reproduced with the first greedy path across the hypercube.
4. *Repeated uniform sampling*: When repeated sampling occurs, it can strengthen the inference around where traits are acquired. For example, comparing the first four traits of Fig. C.9(i) and Fig. C.10(i), we can see that the repeated sampling afforded by 10 repeated trajectories almost completely removes any density for acquisition off the progression.
5. *Non-uniform sampling across the progression*: We consider this assumption in Fig. C.11. When some states are sampled a greater number of times, parameterisations that lead to this state will have a stronger ‘signal’ than those where the observation just occurs once. We illustrate this important effect with several examples. In all cases we consider the complete left-right progression but with the state *s* = 11110000 sampled 100 times more than the others. In Fig. C.11(i) we see this state acts as a ‘gateway’ by removing uncertainty for the acquisition of features present in *s* after *s* is encountered, and removing uncertainty in acquisition of features absent in *s* before *s* is encountered. In Fig. C.11(ii), the right-left progression is also included but with uniform sampling. The non-uniform sampling leads to a much greater representation of the left-right progression. In Fig. C.11(iii), two noisy trajectories are now included (only uniform sampling for the noisy trajectories). As the noise is uniform, acquisitions before *s* still clearly resemble the progression, while features not present in *s* become affected by the noise.

**Fig. C.10:**
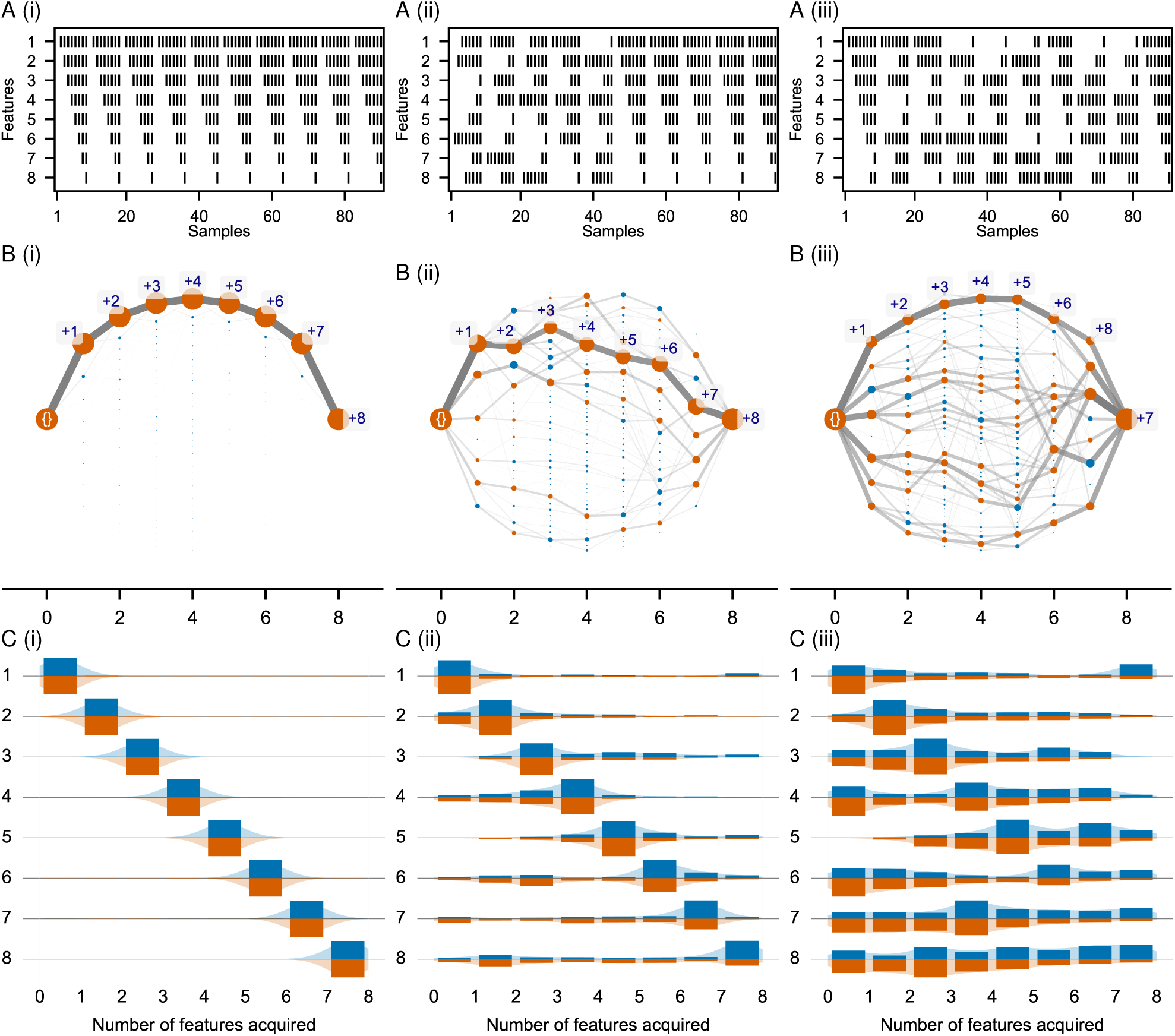
HyperTraPS inference in the presence of noisy samples. (i) One complete progression with ten samples from each state instead of a single sample (*|D*| = 10*L* compared to *|D|* = *L*). (ii) Five out of the ten trajectories part of the dataset involved the features being randomly acquired instead of the left-right progression. (iii) Nine out of the ten trajectories part of the dataset involved the features being randomly acquired instead of the left-right progression. The figure structure mimics that of Fig. C.9. For (i), the hypercube plot and histogram plot shows more tightly defined paths due the ten-fold increase in data supporting the primary pathway, pushing the posterior towards the maximum likelihood parameterisation. In (ii), the introduction of this noise is visible but does not obscure the dominant non-noisy progression from being disambiguated. (iii) For (iii), the introduction of the uniform noise has a significant effect on the nature of paths observed across the hypercube, although even in this case it should be noted the appearance of the first greedy path being almost identical in structure to the non-noisy path structure.

## Appendix D. Detailed comparison of HyperTraPS with alternative Bayes network approach

### Appendix D.1. Comparison for synthetic datasets

In this section we look in more detail at how our main three synthetic datasets and the ovarian dataset can be analysed with HyperTraPS or an alternative approach with Bayesian networks. We use the same three datasets 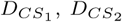 and 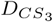 but with *L* = 7 for additional comparability with the ovarian dataset. We perform model fits with HyperTraPS and the Capri model from the Tronco package (Loohuis et al., 2014). Capri aims to output a maximum likelihood DAG relying upon a Suppes-Bayes plausible causation condition for an edge to be permitted between two features in the dataset, creating a Bayes network. Temporal priority and probability raising are key to this approach for feature relationship reconstruction (Loohuis et al., 2014). That is, if the data suggests that *P* (*A*) *> P* (*B*) and 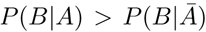, *A* may be considered to be a plausible cause of *B*, allowing temporal and causal relationships between features to be investigated. Regularisation procedures allow for optimal graph structures to be found for a given dataset based on rules for disregarding edges of the feature to feature graph that are less supported by the data.

**Fig. C.11:**
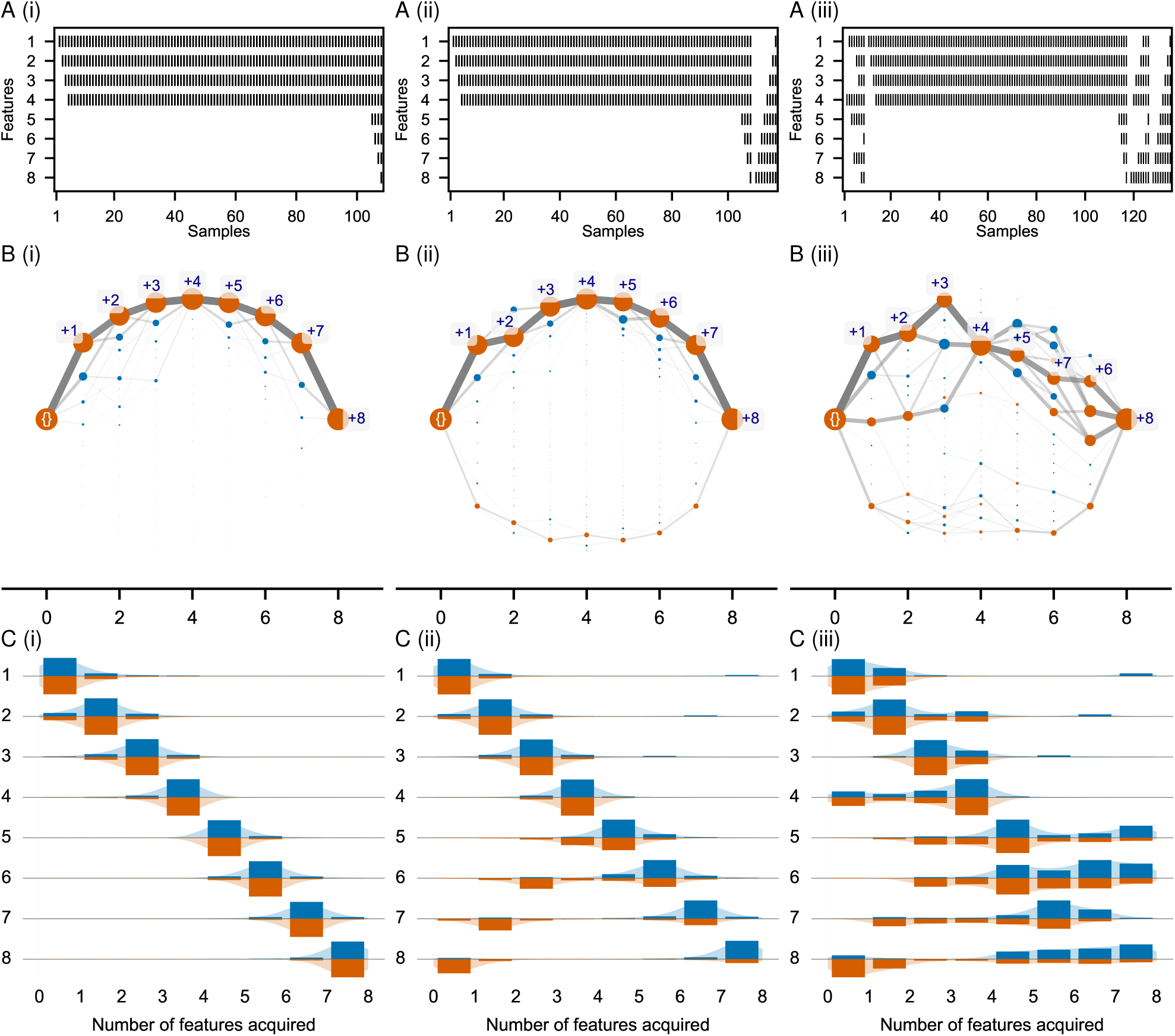
HyperTraPS inference in the presence of non-uniform sampling. In each of (i)-(iii) the state *s* = 11110000 is sampled 100 times more than all other samples. (i) Only the single left-right progression. (ii) The single left-right progression with the non-uniform sampled middle state is present and a second progression with uniform sampling from right-left. (iii) Same as (ii) but a single noisy progression is added in each direction. The figure structure mimics that of Fig. C.9 and C.10. For (i), the oversampled state acts as a gateway with uncertainty remaining in the regions where acquisition occurs before and after the gate. For example, *f*_45_ *≈* 0 in contrast to Fig. C.9(a), while *f*_43_ 0 as for the uniform case. For (ii), where two progressions are present but only the left-right has oversampling in the middle, due to the oversampling in the left-right path there is a large bias towards random walks from 0^*L*^ following this path, as seen by the strength of corresponding path in the hypercube plot. WS2 allows for this to be accounted for illustrating the other pathway more clearly as the simulations ensure the right-left progression is visited. For (iii), noise is now introduced for both progressions. As the noise is uniform, acquisitions before the oversampled state *s* still resemble the dominant progression, while subsequently the noise clearly affects the order of acquisition increasing the uniformity of feature acquisition. The right-left progression becomes difficult to distinguish at all due to a lack of random walks beginning at 0^*L*^ following this progression. However, the ability for the inference to perform random walks that take this weaker and noisy second progression is remarkable as observed by the fact orange states from the data associated with the progression are still encountered.

**Table D.2:**
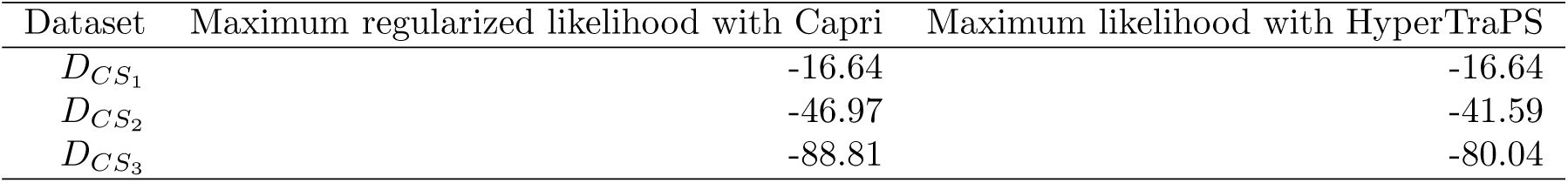
Maximum likelihood values for Capri and HyperTraPS with each sythetic cross-sectional dataset and ovarian CGH dataset. Where there is a single progression (*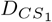*) both models reproduce the same maximum likelihood via the two completely different approaches. Where there is more than a single progression (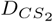 and 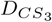), the additional stochastic flexibility available in HyperTraPS parameterisations allows models with larger maximum likelihoods to be recovered without the enforcement of monotonicity.

**Fig. D.12:**
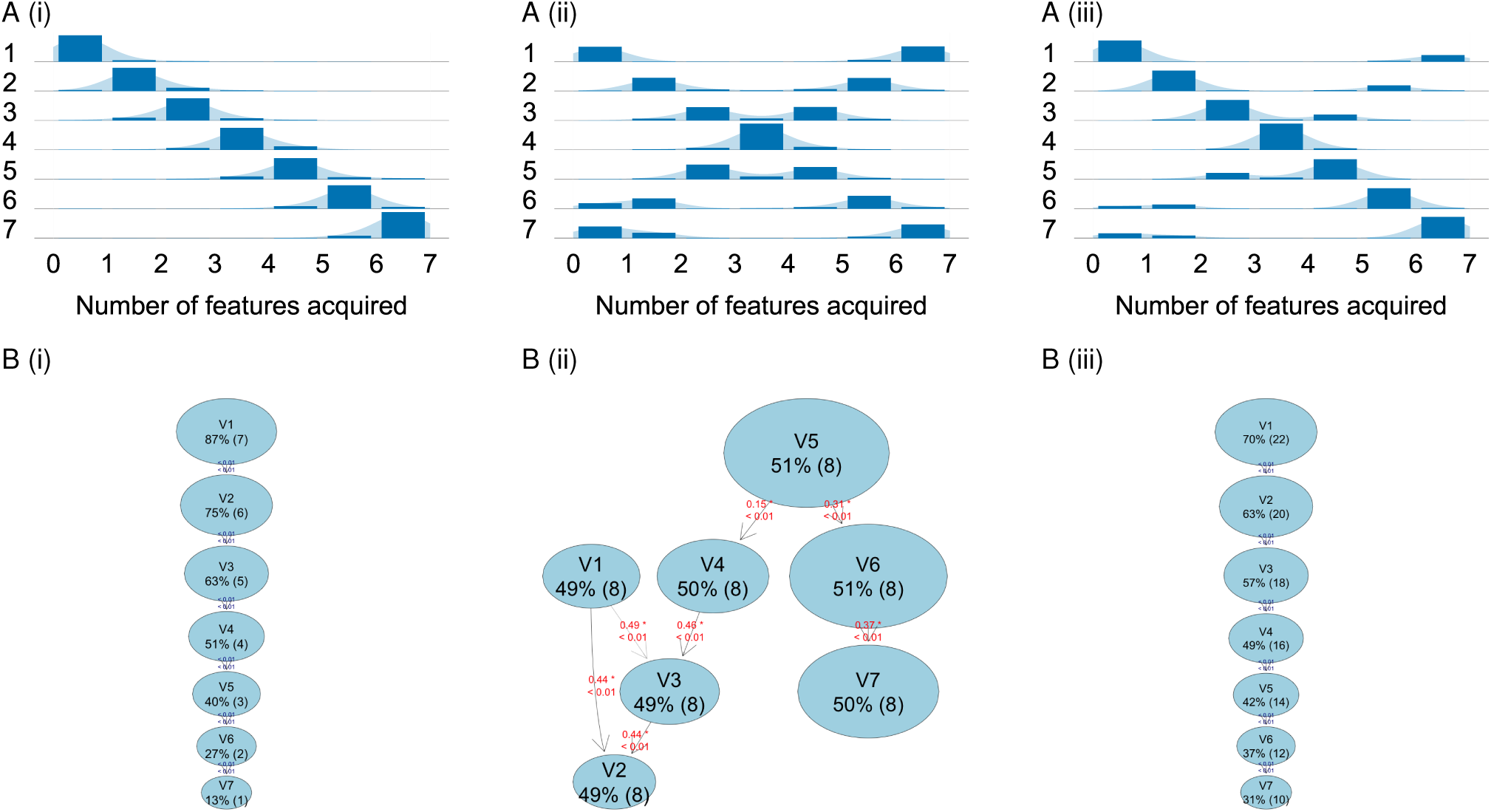
HyperTraPS outputs A(i)-(iii) compared with Capri outputs B(i)-(iv) from the Tronco package (De Sano et al., 2016) applied to 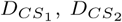 and 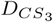. Each edge for Capri has a p-value associated with the significance of the edge for temporal priority and probability raising from bootstrapping of the data. In A(i) and B(i), the single underlying progression in 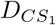 is reproduced significantly in both methods. In A(ii) and B(ii) for 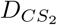 where two progressions (in equal proportion) are present, HyperTraPS distinguishes the two while Capri cannot significantly distinguish due to a lack of temporal priority for all features as they form part of progressions with different orderings. In A(iii) and B(iii), for 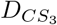, there is a dominant progression which HyperTraPS and Capri identify, while only HyperTraPS identifies the second potential progression, evidenced by the additional density from the bottom left to the top right of the feature histogram.

Table D.2 shows the resulting maximum likelihoods achieved in each model. The likelihoods here are made comparable by incorporating the *P*_emit_(*O, s*_*i*_) for each observed state in the dataset. The same maximum likelihood is recovered for the 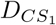 dataset where there is a single temporal progression for features in the data, while greater maximum likelihoods are recovered for 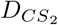 and 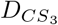 are recovered with HyperTraPS. This provides an illustration of data where HyperTraPS can derive models with greater associated likelihoods than approaches where a single progression of features is an implicit assumption of the method due to the platform’s ability to capture multiple temporal pathways through its stochastic foundations.

In Fig. D.12A(i)-(iii) and D.12B(i)-(iii), the outputs of HyperTraPS compared to Capri illustrate this by the two progressions present in 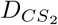 and 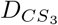 being clearly visible in the HyperTraPS histograms but not detectable with the graphical model reconstructed by Capri.

### Appendix D.2. Comparison for ovarian dataset

We consider the comparison further for the biomedical dataset in the main text for ovarian cancer alterations. The ovarian-cgh data has been previously analysed using Bayesian network approaches in two distinct ways: one using oncogenetic trees with *Oncotrees* (Desper et al., 1999; Szabo and Boucher, 2002) *and the Suppes-Bayes approach discussed here with a slightly different method for pruning edges from the feature to feature graph (the Caprese* algorithm (Loohuis et al., 2014)).

In Fig. D.13A and D.13B, we show the outputs for HyperTraPS (with a the hypercube plot and histogram plots from the main text) and the output of the Capri algorithm (Ramazzotti et al., 2015) respectively as for the synthetic datasets. In this case, HyperTraPS generates significant variation in the order of acquisition of features, as indicated by a lack of a strong mode for several of the features (*5q-* and *4q-* that have quite uniform distributions. In contrast, the Capri algorithm presents a clear relationship of precedence between all the features in a maximum likelihood DAG that is significant both in temporal priority and probability raising (*p <* 0.01 in the plot). Despite the clear difference in outputs from the method, there are still clearly strong apparent similarities such as the frequency of *8q+* over *3q+* as the first genetic acquisition.

**Fig. D.13:**
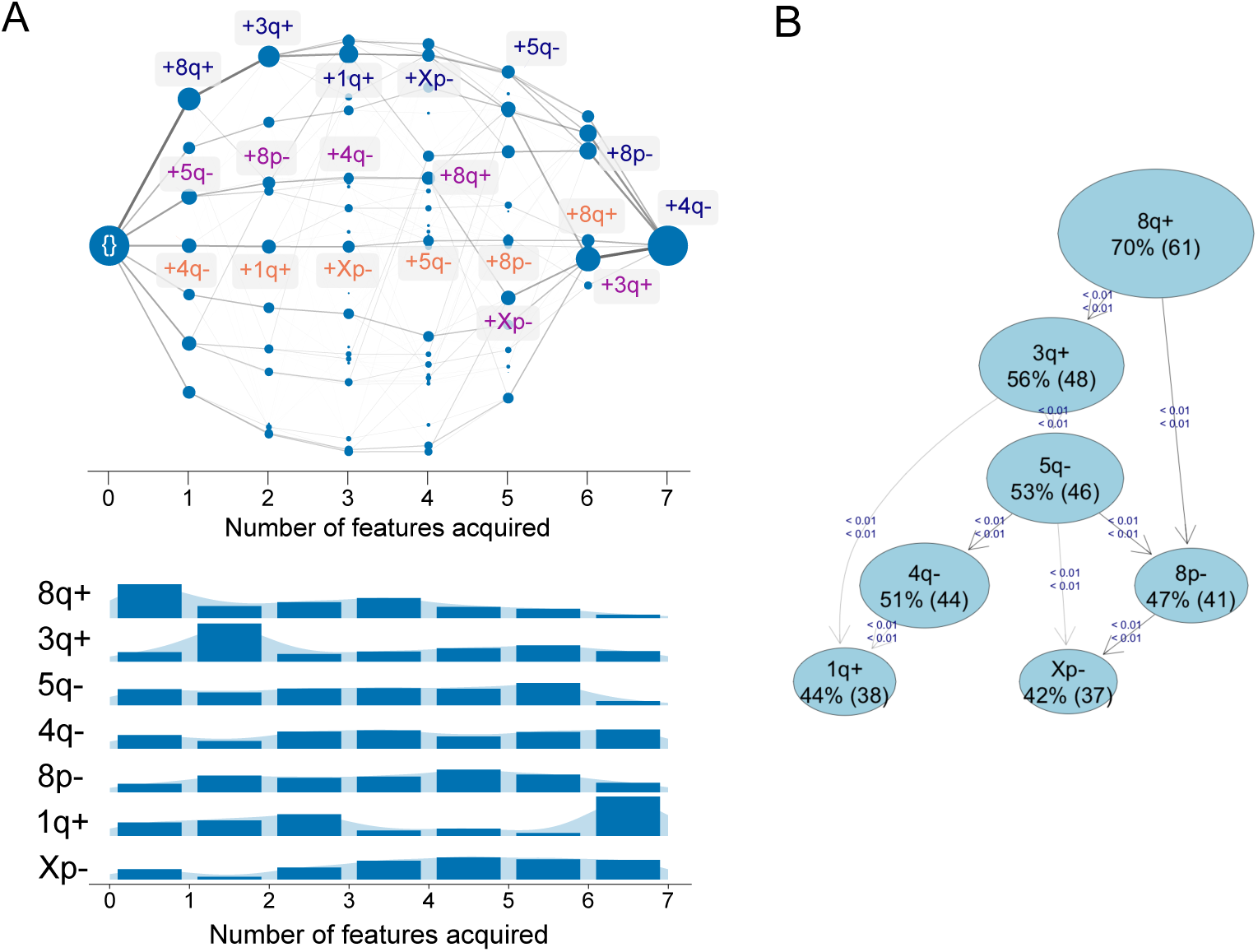
(A) HyperTraPS outputs compared with (B) Capri outputs from the Tronco package (De Sano et al., 2016) applied to the ovarian dataset.

Our aim with this comparison is not to claim one approach is superior to the other, but to highlight that the HyperTraPS framework provides an alternative, inherently probabilistic, methodology for analysing such data that generates different model outputs that may provide different insights towards understanding such systems, as discussed in the Main Text. As such, the methods can be considered as complementary: HyperTraPS can be utilised to highlight the possibility of multiple underlying progressions with different temporal priorities for the features, while the Bayesian network approaches can attempt to precisely reconstruct the feature relationships through imposing more restrictive assumptions.

## Appendix E. Additional interpretation of findings for tuberculosis dataset

Additional comparisons can be made between the inferred order of polymorphism acquisition in Fig. 4 and Fig. B.8B and the findings of by Casali et al. (2014). Of the *L* = 19 features used for the analysis, we pick a subset here that provide interesting discussion points with regard to co-associations discussed by the authors. These points demonstrate the ability of HyperTraPS to provide quantitative support for existing hypotheses, and to suggest new avenues of mechanistic research, in complex biological systems.

- *Drug-resistance and fitness compensatory mutations*: Of the *L* = 19 features, the first 16 correspond to the drug-resistant polymorphisms within genes or in the promoter regions. The last three (*rpoA, rpoB* and *rpoC* are nonsynonymous SNPs within RNA polymerase genes. The authors considered the occurrence of compensatory mutations in *rpoA* and *rpoC* in response to drug-resistance polymorphism in *rpoB*. WS1 reveals an acquisition ordering with *rpoB* being acquired prior to *rpoC*, suggesting a compensatory effect follows drug-resistance mutations in this case.
- *Mutations arising due to changes in environment*: Casali et al. (2014) find a co-association of *KatG* with *inhA* discussed in the original data, with the the *inhA* mutation arising potentially from the presence of *KatG* polymorphisms in response to specific treatments. This temporal priority of *KatG* followed by *PinhA* mutations are consistent with the HyperTraPS analysis depicted.
- *Genetic sites particularly associated with adaptive selection*: Highly polymorphic genes conferring resistance are known to be *embB, pncA, ethA* (Casali et al., 2014). Interestingly these polymorphisms occur at a wide range of orderings within the inferred orderings, illustrative of their flexibility and why they may be particularly polymorphic – they can play different roles in different progressions.

**Fig. F.14:**
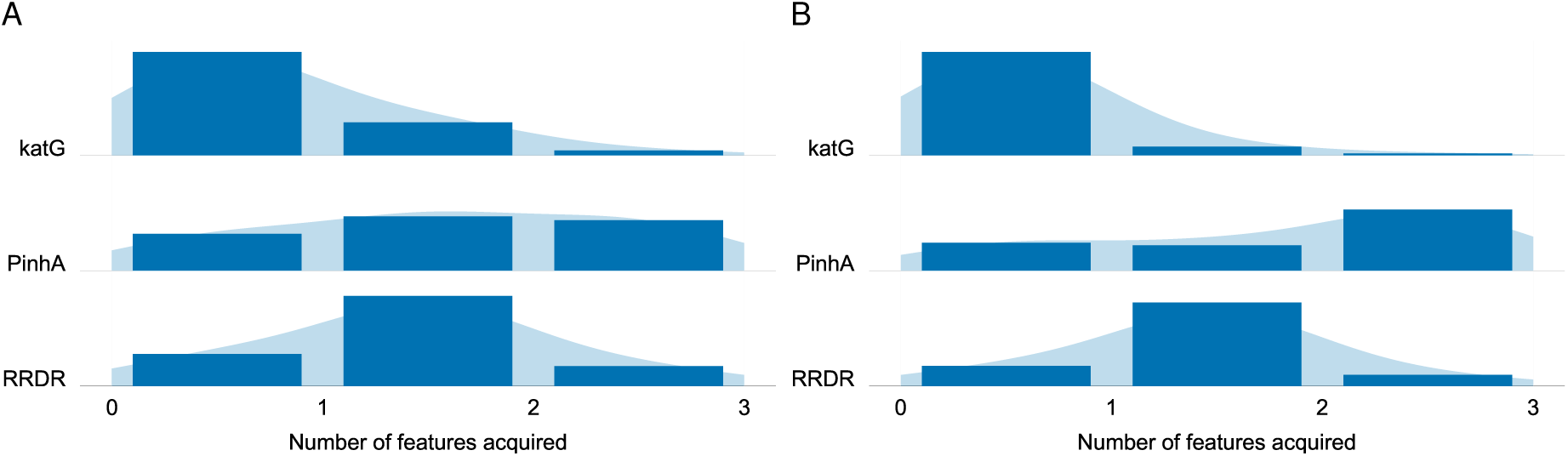
*Comparison of the tuberculosis dataset analysed with both HyperTraPS (A) and Simmap (B) on the restricted, tractable set of genetic sites: KatG, PinhA* and *RRDR*.
- *Transmissibility of drug-resistance*: With respect to transimissibility Casali et al. (2014) suggest that *KatG* is prior to RRDR which is supported in the top two greedy paths highlighted in the hypercube plot in the main text Fig. 4.

## Appendix F. Comparison of HyperTraPS with Simmap for tuberculosis dataset

Here we make a direct comparison of the order in which mutations are acquired with Simmap, which takes the form of a continuous time Markov model with mater equation approach to acquiring characters that belong to leaves on a phylogeny. This approach runs into computational issues when the number of states under evolution grows large (only tractable in short run times for the tuberculosis up to *L≈*5). This is in contrast to HyperTraPS which can handle the full *L* = 19 traits.

As an illustration of compatibility with this alternative approach, we restrict the tuberculosis dataset to *L* = 3 features (*KatG, PinhA* and *RRDR*) with the full set of isolates and enforce single irreversible acquisitions as transitions wuthin the Simmap model in order to make direct comparisons with HyperTraPS. In Fig. F.14A, we show the output for the density of order of acquisition from simulated rate matrices outputted by Simmap with the hypercubic restriction imposed and irreversibility. Alongside in Fig. F.14 we show the result for WS2 with HyperTraPS (as the transitions performed with Simmap are to the sample data and do not fully acquire all features as is the case with WS1). The plots are in close agreement, providing good validation that HyperTraPS generates results consistent with current platforms.

